# Modeling Breast Cancer Proliferation, Drug Synergies, and Alternating Therapies

**DOI:** 10.1101/2022.09.20.508795

**Authors:** Wei He, Diane M. Demas, Ayesha N. Shajahan-Haq, William T. Baumann

## Abstract

Estrogen receptor positive (ER+) breast cancer is responsive to a number of targeted therapies used clinically. Unfortunately, the continuous application of targeted therapy often results in resistance. Mathematical modeling of the dynamics of cancer cell drug responses can help find better therapies that not only hold proliferation in check but also potentially stave off resistance. Toward this end, we developed a mathematical model that can simulate various mono, combination and alternating therapies for ER+ breast cancer cells at different doses over long time scales. The model is used to look for optimal drug combinations and predicts a significant synergism between Cdk4/6 inhibitors in combination with the anti-estrogen fulvestrant, which may help explain the clinical success of adding CDK4/6 inhibitors to anti-estrogen therapy. Lastly, the model is used to optimize an alternating treatment protocol that works as well as monotherapy while using less total drug dose.

## INTRODUCTION

Metastatic breast cancer remains an incurable disease and it is estimated that 43,250 women and men will die from breast cancer this year (Siegel et al., 2022). The most common type of breast cancer, estrogen receptor positive (ER+), which is present in approximately 70% of all breast cancers (Özdemir et al., 2018), has targeted therapies that have dramatically improved long-term survival rates (Chia et al., 2010; Tremont et al., 2017; Xi et al., 2020). However, the continuous application of these drugs can ultimately lead to drug resistance and recurrence (Gururaj et el., 2006; Zhou et al.,2007; Osborne et al., 2011; Lei et al., 2019; Xi et al., 2020). The resistance mechanisms are varied and include epigenetic changes, gene mutation, amplification and deletion (Musgrove et al., 2009; Sharma et al., 2010; Tilghman et al., 2013; Ma et al., 2015; Herrera-Abreu et al., 2016). While targeted therapies are important methods for breast cancer treatment, eventually cancer cells become resistant and proliferate again, which makes the advantage of targeted therapies only temporary for many patients.

Constant application of one drug regimen over time may not be optimal, but to move beyond this approach requires addressing a number of critical questions such as (1) how long should a given therapy be applied, (2) what should the next therapy be, and (3) in any given therapy interval, what is the best combination of drugs to apply? These questions are difficult to answer experimentally, even *in vitro*, as long time scales are involved and there are a huge number of possible solutions to explore. Systematic application of an experimentally-calibrated mathematical model that integrates molecular cell biology and drug pharmacology can help us investigate better treatment regimens in terms of drug choice, combinations, dosing and scheduling (Lalonde et al., 2007; Visser et al., 2014; Chakrabarti et al., 2017; Zhang et al., 2017). In this work, we take a step towards answering these questions in a common ER+ breast cancer cell line, MCF7, by using a combination of mathematical modeling and experimental investigations. Previously, we developed a mechanistic mathematical model based on key interactions between ER signaling and the cell cycle (He et al., 2020). This model was calibrated using protein and proliferation data from 7-day time courses of MCF7 cells growing under basal conditions or responding to standard clinical drugs in ER+ breast cancer: (1) estrogen deprivation (–E2), a surrogate for an aromatase inhibitor that lowers the estradiol (E2) level by inhibiting aromatase (Ma et al., 2015; Seruga et al., 2019); (2) ICI 182 780 (ICI; Faslodex/fulvestrant), a proteasome-dependent ER degrader (Wittmann et al., 2007); or (3) palbociclib, a Cdk4/6 inhibitor (Xi et al., 2020). To address questions regarding synergies, longer time scales, and alternating treatments, more experimental data is required to either validate the initial model or show where extensions to the model are required.

In this study, we extend the model to handle a range of doses of ICI or palbociclib and to more accurately predict proliferation over longer time scales and in cases where drugs are changed periodically. Key extensions involve the accumulation of cyclinD1 and the long-term slowdown in growth rate in response to continuous palbociclib treatment. We use the resulting model to explore synergistic drug combinations and find a combination that allows a significant reduction in overall drug dose compared to monotreatment. The model is also used to optimize an alternating treatment intended to delay the development of resistance and finds a protocol that has the same proliferation as monotherapy while using a significantly lower total drug dose.

## RESULTS

Mathematical models with many parameters and limited experimental calibration data, as is the case here, have many possible parameter sets that do a reasonable job of fitting the data. Therefore, in addition to the best-fit parameter set, we created a cohort of 199 additional parameter sets that increase the fitting cost less than about 25% over the optimal (see Star Methods). When plotting our results, we plot the best parameter set as a solid line and use shading to indicate the range of results from simulating the entire cohort, to give some idea of how much the training data constrains the simulation results. We also note that we only write that the model “predicts” something if the model simulation is being compared to experimental data on which it was not trained. In all other cases the plots show the simulations recapitulating the training data (see Table S1). All simulation results use the final version of the model.

### Simulating Proliferation under Constant Therapy

Based on the effect of estrogen signaling and Cdk4/6 inhibition on the G1-S transition of the cell cycle (Musgrove, et al., 2009; Lynce et al., 2018), we built a mechanistic mathematical model using ordinary differential equations (ODEs). The biological interactions we considered are based on known mechanisms from the literature and are shown in Figure 1A. The details and references for each numbered interaction are provided in STAR Methods. To create the ODE model, we modified and simplified the interactions shown in Figure 1A. In particular, we used the RB1-pp (hyperphosphorylated form of retinoblastoma protein (RB1)) level to reflect the transcriptional activity of E2F and associated the RB1-pp level with proliferation. The model structure is shown in Figure 1B and the explanations of the modifications and simplifications are provided in STAR Methods.

**Figure 1.**
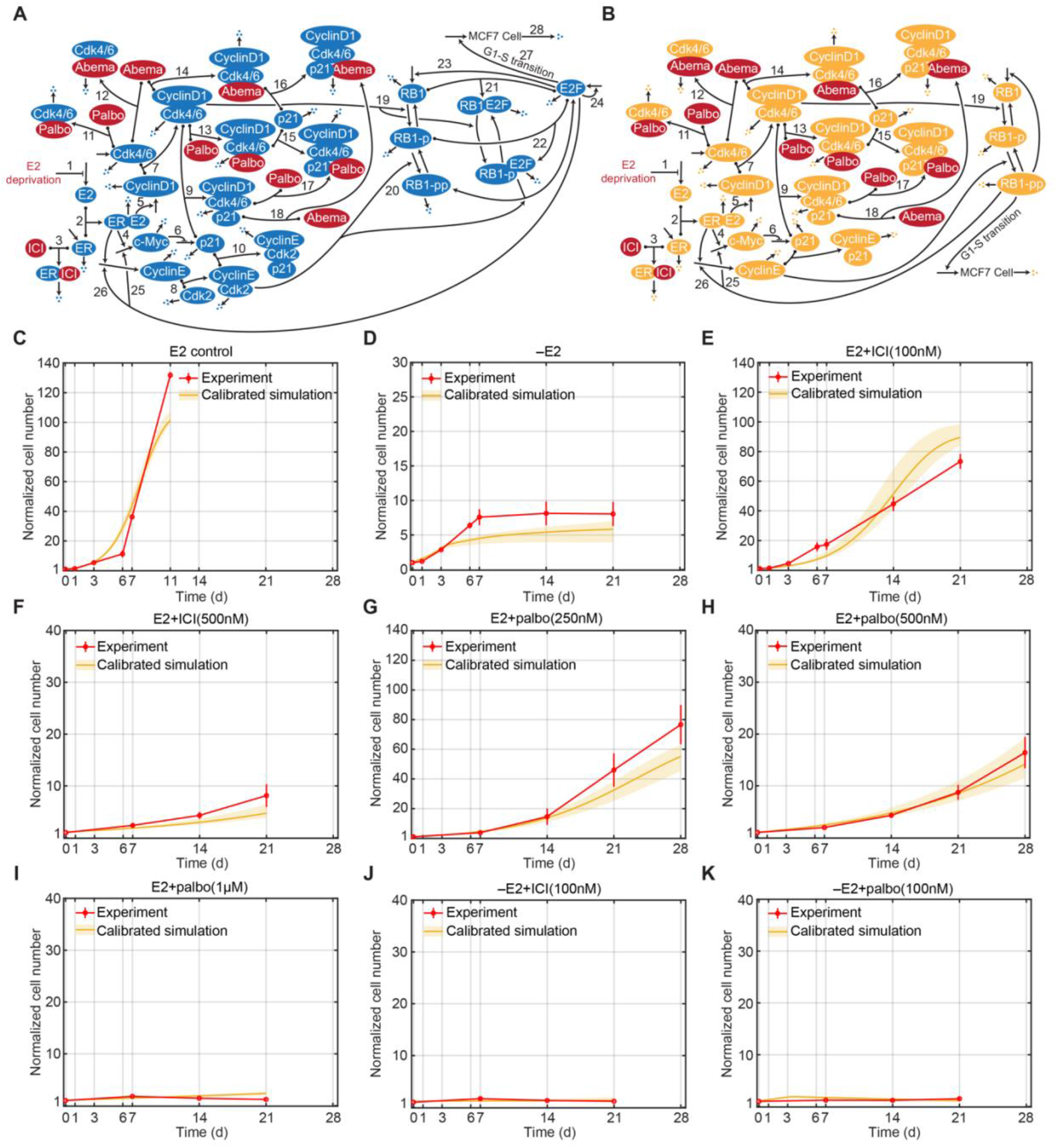
Signaling Diagram of the Biological Mechanism, Model Structure and Model Calibrations. (A) Detailed reactions of the biological mechanism related to estrogen signaling and Cdk4/6 inhibition. Reversible binding reactions are represented by dots on the components and an arrow to the complex. Three dots represent degradation of a protein or the death of a cell. Arrows pointing from blank space to a protein or MCF7 cell represent production of the protein or proliferation of the cell. Arrows pointing from one protein to another protein represent phosphorylation or dephosphorylation of the protein. Lines pointing to other lines represent enhancement (arrow) or inhibition (blunt head) of the reactions. Treatments are colored red. The numbered biological mechanism consisting of the following process: 1. –E2 decreases estrogen; 2. E2 binds to ER; 3. ICI binds to ER; 3. E2:ER increases transcription of c-Myc; 5. E2:ER increases transcription of cyclinD1; 6. c-Myc inhibits transcription of p21; 7. CyclinD1 binds to Cdk4/6; 8. CyclinE binds to Cdk2; 9. p21 binds to cyclinD1:Cdk4/6; 10. p21 binds to cyclinE:Cdk2; 11. Palbociclib binds to Cdk4/6; 12. Abemaciclib binds to Cdk4/6; 13. Palbociclib binds to cyclinD1:Cdk4/6; 14. Abemaciclib binds to cyclinD1:Cdk4/6; 15. p21 binds to cyclinD1:Cdk4/6:palbociclib; 16. p21 binds to cyclinD1:Cdk4/6:abemaciclib; 17. Palbociclib binds to cyclinD1:Cdk4/6:p21; 18. Abemaciclib binds to cyclinD1:Cdk4/6:p21; 19. CyclinD1:Cdk4/6 phosphorylates RB1; 20. CyclinE:Cdk2 phosphorylates RB1-p; 21. RB1 binds to E2F; 22. RB1-p binds to E2F; 23. E2F up-regulates RB1; 24. E2F up-regulates itself; 25. E2F up-regulates c-Myc; 26. E2F up-regulates cyclinE; 27. E2F drives the G1-S cell cycle transition and proliferation; 28. Cell death. (STAR Methods). (B) Structure of the mathematical model, a simplified version of the biological mechanism in (A). (C) Model calibration to experimental data (mean ± s.e., n=3) in E2 control condition. The experimental data are shown in red and the calibration simulation results are shown in yellow (solid line represents the lowest cost value simulation and the shaded regions contains the central 98% of the cohort simulations). (D) Model calibration to experimental data (mean ± s.e., n=3) in –E2 condition. (E) Model calibration to experimental data (mean ± s.e., n=3) in E2+ICI(100nM) condition. (F) Model calibration to experimental data (mean ± s.e., n=3) in E2+ICI(500nM) condition. (G) Model calibration to experimental data (mean ± s.e., n=3) in E2+palbo(250nM) condition. (H) Model calibration to experimental data (mean ± s.e., n=3) in E2+palbo(500nM) condition. (I) Model calibration to experimental data (mean ± s.e., n=3) in E2+palbo(1μM) condition. (J) Model calibration to experimental data (mean ± s.e., n=3) in –E2+ICI(100nM) condition. (K) Model calibration to experimental data (mean ± s.e., n=3) in E2+palbo(100nM) condition.

Figures 1C–1K compare the model simulation results of 21 or 28-day proliferation to experimental results for numerous treatments. Figure 1C shows the cell proliferation in the E2 control condition (E2 control), which is much faster than that in other mono and combination treatment conditions. The E2 control experiment was stopped early, at day 11, due to confluence. In the E2 deprivation (–E2) condition, shown in Figure 1D, cells proliferate during the first 7 days and then essentially stop. This effect is captured by the model by adding the dynamics of E2 concentration to the model, where the E2 concentration decreases with each medium change, increasingly depriving the ER of its ligand (see STAR Methods and Figures S6A-B). The –E2 experiment illustrates how cell proliferation over longer time scales can be qualitatively different from that over short time scales, so a mathematical model calibrated on short time scale experiments may not be useful for simulations on a longer time scale, hence the necessity of long time scale experimental data. Figures 1E and Figure 1F show the decrease in proliferation, due to increased ER degradation, as the dose of ICI treatment (E2+ICI) increases from 100nM to 500nM. Figures 1G–1I show the decreasing proliferation, due to increased Cdk4/6 inhibition, as the dose of palbociclib treatment (E2+palbo) increases from 250nM, to 500nM, to 1μM. After showing that the model is capable of simulating –E2, ICI and palbociclib monotreatments, Figures 1J and Figure 1K show the model simulation results for two combination treatments, –E2 plus 100nM palbociclib (–E2+palbo) and –E2 plus 100nM ICI (–E2+ICI). Not surprisingly, the combination treatments provide greater effect than either monotreatment by itself. The combination of –E2 and ICI treatments reduces supply of both E2 and ER, causing a larger decrease in the normalized cell number. The combination of –E2 and palbociclib inhibits Cdk4/6:cyclinD1 kinase activity by both reducing the cyclinD1 level and inactivating Cdk4/6, which also causes a larger reduction of proliferation. In addition to the cell number, the model can also capture the protein level changes under –E2 and E2+ICI(500nM) treatments, which were measured in our previous work (He et al., 2020, see Figures S1 and S2).

### Adding a New Drug to the Model

An advantage of a mechanistic mathematical model is that it is straightforward to add a different drug that affects the signaling pathway in the model without requiring extensive experimentation. As the model already captures the mechanism driving changes due to the Cdk4/6 inhibitor palbociclib, adding a different Cdk4/6 inhibitor should require only an update of the small number of parameter values related to Cdk4/6 inhibition. We illustrated this by incorporating one of the other Cdk4/6 inhibitors, abemaciclib (LY2835219), into the model. Abemaciclib is a 2-anilino-2, 4-pyrimidine-[5-benzimidazole] derivative (Roskoski, 2016). Unlike palbociclib, it has been reported to be effective as a single-agent (Dickler et al., 2017; O’Brien et al., 2018; Patnaik et al., 2016). It can inhibit cyclinD1:Cdk4 and cyclinD1:Cdk6 kinase activities at low nanomolar concentration (Gebbia et al., 2020). While at higher micromolar concentrations Abemaciclib has been shown to attack other targets (Knudsen et al., 2017; Cousins et al., 2018; Hafner et al., 2019), we have focused on Cdk4/6 as the most relevant target at the concentrations we consider.

We added the key binding parameters between abemaciclib and Cdk4/6 or Cdk4/6 complexes to the model. In addition to cell number, we measured c-Myc and RB1-pp, two proteins in the model critical to proliferation, to help calibrate the binding parameters associated with abemaciclib. Figures 2A and 2B show that the model can fit the experimental proliferation results for the 300nM and 500nM abemaciclib treatments (E2+abema), respectively. Figures 2C–2D show the protein level changes for c-Myc and RB1-pp in response to 500nM abemaciclib. As expected, abemaciclib inhibits Cdk4/6 activity and decreases the RB1-pp level, which in turn, leads to decreased transcription of c-Myc causing the c-Myc protein level to decrease. Because the mathematical model already captured the mechanism of Cdk4/6 inhibition, it was possible to add another inhibitor of cyclinD1:Cdk4/6 kinase activity without needing to perturb the other signaling pathways.

**Figure 2.**
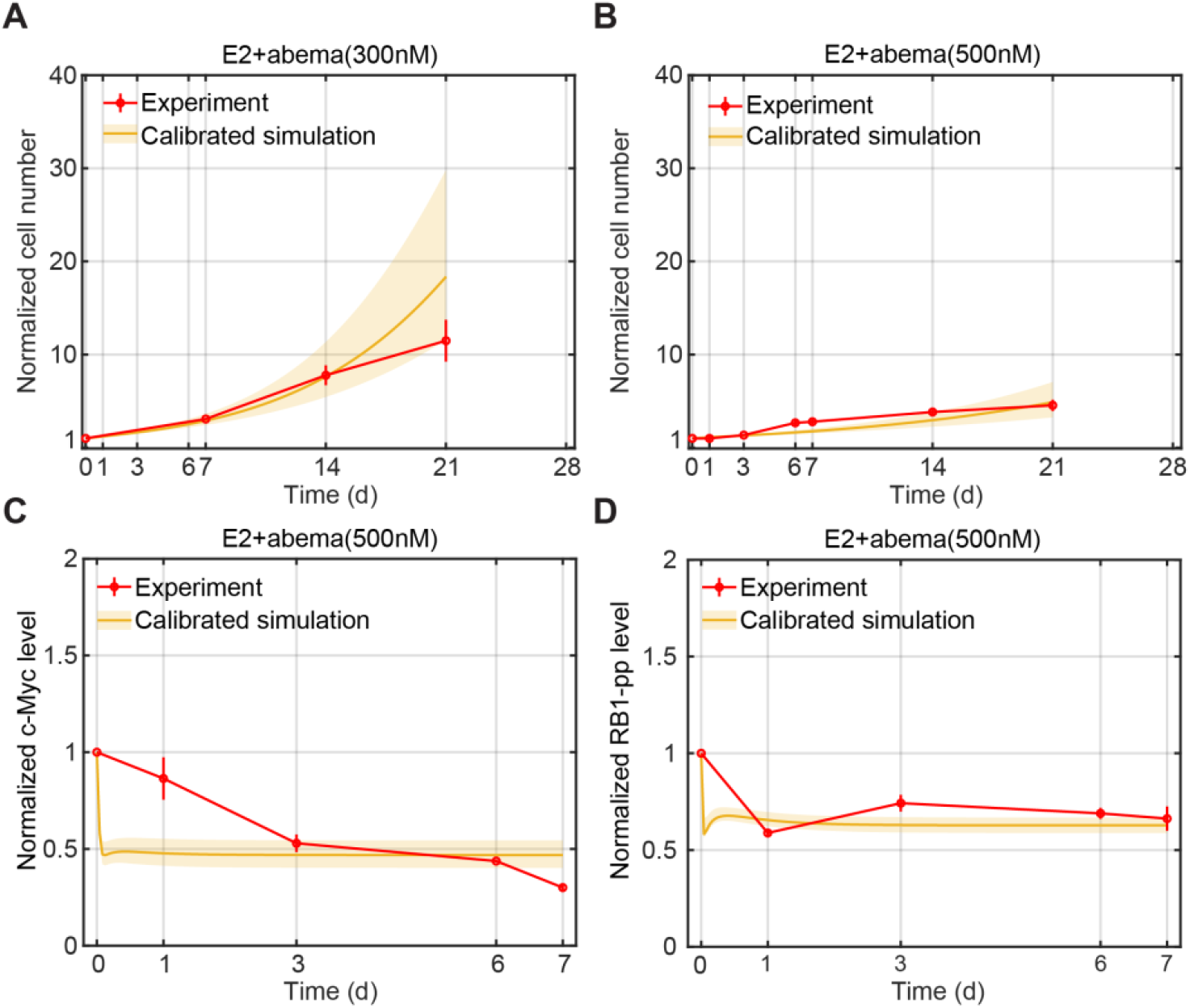
Model Calibration Simulations Compared to Experimental Data for Abemaciclib Treatments. (A) Model calibration of normalized cell number to experimental data (mean ± s.e., n=3) in E2+abema(300nM) condition. The experimental data are shown in red and the calibration simulation results are shown in yellow (solid line represents the lowest cost value simulation and the shaded regions contains the central 98% of the cohort simulations). (B) Model calibration of normalized cell number to experimental data (mean ± s.e., n=3) in E2+abema(500nM) condition. (C). Model calibration of normalized c-Myc level to experimental data (mean ± s.e., n=3) in E2+abema(500nM) condition. (D). Model calibration of normalized RB1-pp level to experimental data (mean ± s.e., n=3) in E2+abema(500nM) condition.

### Simulating Alternating Treatment Involving Estrogen Deprivation

The resistance that develops to continuously applied mono or combination drug therapy represents a significant impediment to successful treatment and we hypothesize that an alternating application of various treatments in a repeating cycle may provide a means of delaying or preventing resistance. Researchers have shown that cancer cell populations can display a transient, reversible, drug-tolerant state to protect the cell from eradication (Sharma et al., 2010; Smith et al., 2016). Therefore, alternating among various drugs may reverse a tolerant state to a given drug back to a sensitive state during the application of a different drug and thereby delay or prevent the development of resistance. Before testing whether alternating treatment can indeed delay the development of resistance, and with an eye toward using the model to design alternating therapies, we first show the model’s capability to simulate proliferation changes in response to alternating therapies.

Figures 3A and 3B show the model simulation results and experimental measurements of two alternating treatments, palbociclib alternating with –E2 and palbociclib alternating with ICI. The duration of each treatment is 7 days and the total treatment period is 28 days. Figure 3A shows E2+palbo(250nM) alternating with –E2. We can see that the model simulation is consistent with the experiment result and the cells proliferate about 90-fold in 4 weeks. This growth increase is larger than we initially expected based on the monotreatment data from Figure 1G and 1D, where cells proliferated about 80-fold under E2+palbo(250nM) monotreatment and proliferation essentially stopped under –E2 after 1 week. The reason for the larger increase is the dynamics of the E2 concentration. The palbociclib treatment has E2 in the medium, which is absorbed by the cells, so when the medium is changed to the –E2 condition the cellular E2 diffuses back into the medium and the resulting concentration is sufficient to drive proliferation (Figure S6A). This palbociclib and –E2 alternating experiment confirms the necessity of incorporating E2 dynamics in alternating treatments involving deprivation. Figure 3B shows palbociclib(500nM) alternating with ICI(500nM). We can see that the model simulation is consistent with the experimental result and the cells proliferate about 27-fold in 4 weeks. We conclude that when alternating palbociclib with an endocrine treatment in cell culture, ICI is a better choice than –E2 in terms of controlling the proliferation.

**Figure 3.**
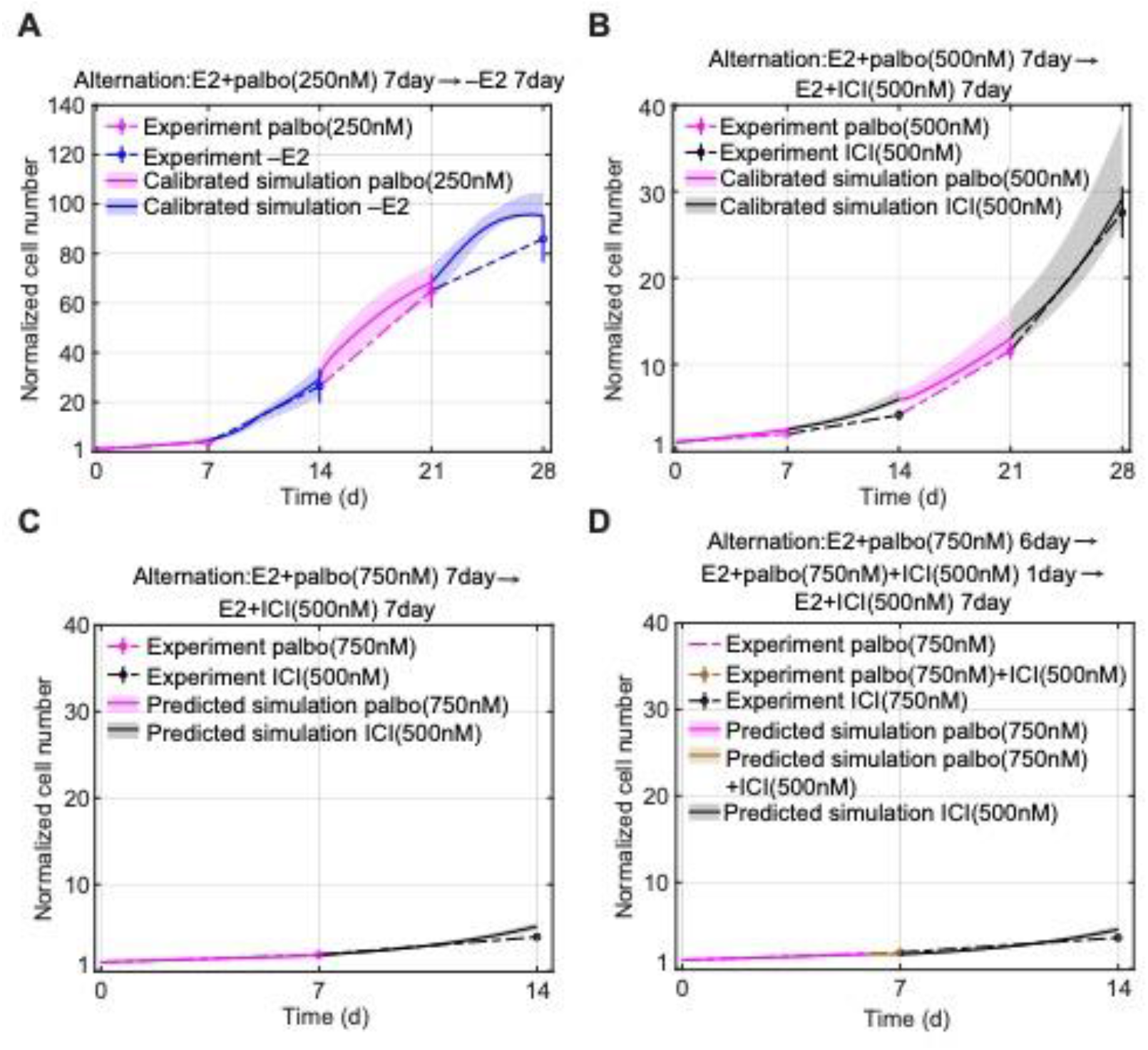
Model Calibration and Prediction Simulations of Normalized Cell Number Compared to Experimental Data for Alternating Treatments. (A) Model calibration to experimental data (mean ± s.e., n=3) of E2+palbo(250nM) alternating with –E2 treatment. The experimental data is linked by dashed lines. The E2+palbo(250nM) treatment is shown in purple and the –E2 condition in blue. The calibration simulation results are shown in the same colors as the experimental data with the solid line representing the lowest cost value simulation and the shaded regions containing the central 98% of the cohort simulations. (B) Model calibration to experimental data (mean ± s.e., n=3) of E2+palbo(500nM) alternating with E2+ICI(500nM) treatment. E2+palbo(500nM) treatment is shown in purple and E2+ICI(500nM) in black. (C) Model prediction of experimental data (mean ± s.e., n=3) for E2+palbo(750nM) alternating with E2+ICI(500nM). E2+palbo(750nM) treatment is shown in purple and E2+ICI(500nM) in black. The treatment started with E2+palbo(750nM) with 7days then altered to E2+ICI(500nM) with 7days. (D) Model prediction of experimental data (mean ± s.e., n=3) for E2+palbo(750nM) altering with E2+palbo(750nM)+ICI(500nM) and E2+ICI(500nM) treatment. E2+palbo(750nM) condition is shown in purple, E2+palbo(750nM)+ICI(500nM) in brown and E2+ICI(500nM) in black. The treatment started with E2+palbo(750nM) for 6days, then changed to E2+palbo(750nM)+ICI(500nM) for 1day and then changed to E2+ICI(500nM) for 7 days.

After showing that the model can simulate these two alternating treatments, we check whether the model can predict the effects of other alternating treatments. Figures 3C and 3D show the model prediction and experimental measurements of the normalized cell numbers under two alternating treatments. The first alternating treatment shown in Figure 3C is palbociclib(750nM) alternating with ICI(500nM). The duration of each treatment is 7 days and the total treatment period is 14 days. The second alternating treatment shown in Figure 3D is palbociclib(750nM) for 6 days, followed palbociclib(750nM) plus ICI(500nM) for 1 day, followed by ICI(500nM) for 7 days. The difference between the first and second alternating treatment is that the second treatment adds a 1 day overlap of palbociclib(750nM) plus ICI(500nM) treatments. Therefore, as shown in Figure 3D, the total experimental proliferation of the second alternating treatment is slightly smaller than the first alternating treatment due to this 1 day combination treatment which has a stronger inhibition effect compared with monotreatment (mean values are 3.9 and 3.5, respectively). The model prediction for the second alternating treatment is also smaller than the prediction for the first alternating treatment as well (mean values are 5.2 and 4.6, respectively).

### Modeling Palbociclib/ICI Alternating Therapy Over Longer Time Scales

One goal of this study is to test whether an alternating treatment can indeed impact the development of resistance. In patients, resistance to Cdk4/6 inhibitors can occur within months (Shah et al., 2018; Pandey et al., 2019), compared with endocrine resistance that may take years to fully develop (Song et al., 2001; Chan et al., 2002; Song et al., 2005). Based on this observation, we decided to test whether an alternating treatment of ICI and palbociclib can affect the development of resistance to palbociclib. A 10 week experiment was conducted where palbociclib was alternated with ICI at weekly intervals. Monotreatment with palbociclib or ICI were included as controls. Based on the results from Figures 1C–1K and Figure 3, we chose the palbociclib and ICI drug doses to be 750nM as our model at that time indicated this dose would cause relatively low proliferation for the controls as well as the alternating treatment and enable the experiment to run without replating. Figure 4A shows the experimental and simulated cell proliferation results for the 10 week protocol. Two major results from the experiment that required adjustments to the initial model were: (1) cells undergoing palbociclib monotreatment grew more slowly as time went on and (2) cell proliferation was much greater than expected in cells that received the alternating drugs, forcing a replating at week 5 to avoid confluence.

**Figure 4.**
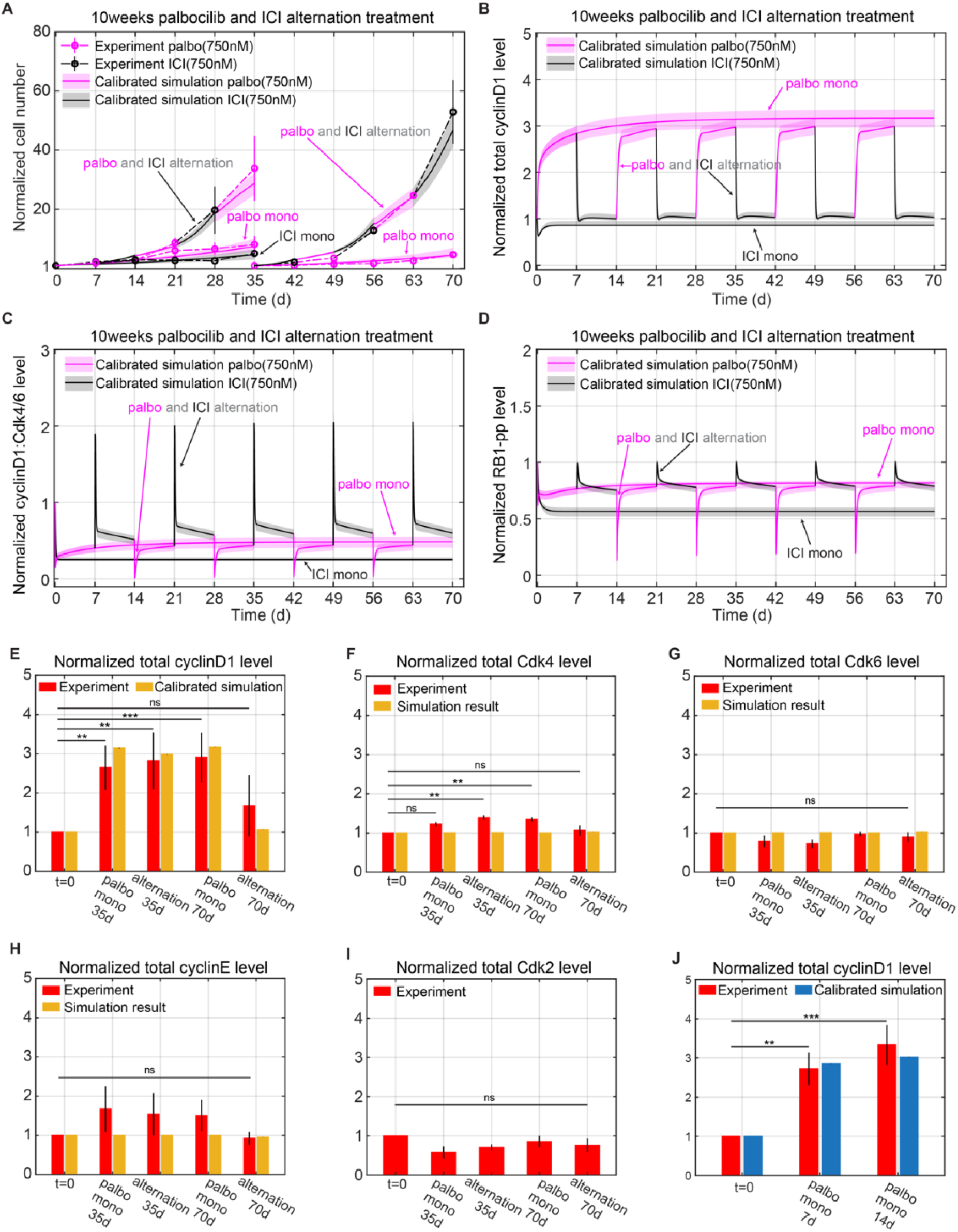
Model Simulations of Normalized Cell Number and Protein Level Changes Compared to Experimental Data for Long Time Mono and Alternating Treatments. (A) Model calibration to experimental data (mean ± s.e., n=3) for E2+palbo(750nM), E2+ICI(750nM), and E2+palbo(750nM) alternating with E2+ICI(750nM) treatments. The experimental data are linked by dashed lines. In both the mono and alternating treatments, the E2+palbo(750nM) condition is shown in purple and the E2+ICI(750nM) condition in black. In the alternating treatment, each treatment period is 7days, starting with E2+palbo(750nM). MCF7 cells are re-plated at 35days in the E2+palbo(750nM) mono and alternating treatments. The normalized cell number from 35 to 70 days is relative to the number plated at 35days. The calibration simulation results are shown in same color as the experimental data with the solid line representing the lowest cost value simulation and the shaded regions containing the central 98% of the cohort simulations. (B) Model simulation of normalized total cyclinD1 level changes in the mono and alternating treatments shown in (A). (C) Model simulation of normalized cyclinD1:Cdk4/6 level changes in the mono and alternating treatments shown in (A). (D) Model simulation of normalized RB1-pp levels changes in the mono and alternating treatments shown in (A). (E) Bar plot of model simulation for total cyclinD1 level compared to experimental data (mean ± s.e., n=3) in E2+palbo(750nM) and E2+palbo(750nM) alternating with E2+ICI(750nM) treatments shown in (A). Total cyclinD1 levels are measured at 35 days and 70 days. The simulation results shown in yellow are the average results from all cohort simulations. Statistical testing was performed by one-way ANOVA (ns: non-significant; *: p <0.05; **: p ≤ 0.01; ***: p ≤ 0.001; ****: p ≤ 0.0001). (F) Bar plot of model simulation and experimental results (mean ± s.e., n=3) for total Cdk4 level changes in E2+palbo(750nM) and E2+palbo(750nM) alternating with E2+ICI(750nM) treatments shown in (A). (G) Bar plot of model simulation and experimental results (mean ± s.e., n=3) for total Cdk6 level changes in E2+palbo(750nM) and E2+palbo(750nM) alternating with E2+ICI(750nM) treatments shown in (A). (H) Bar plot of model simulation and experimental results (mean ± s.e., n=3) for total cyclinE level changes in E2+palbo(750nM) and E2+palbo(750nM) alternating with E2+ICI(750nM) treatments shown in (A). (I) Bar plot of experimental results (mean ± s.e., n=3) for total Cdk2 level changes in E2+palbo(750nM) and E2+palbo(750nM) altering with E2+ICI(750nM) treatments shown in (A). (J) Bar plot of model calibration for total cyclinD1 level changes to experimental data (mean ± s.e., n=3) in E2+palbo(750nM) treatment. Total cyclinD1 levels are measured at 7days and 14days. The statistical testing is the same as (B). The simulation results are average results from all the cohort simulations.

The first 5 weeks of palbociclib monotreatment results in an 8.1-fold increase in cell number while the second 5 weeks results in a 4.6-fold increase (Figure S4). In order to account for this effect, a phenomenological variable, *respropalbo* (number 25 in Table 1), was added to the model to gradually slow down growth in response to long-term palbociclib treatment. This inhibition effect increases gradually when palbociclib is being applied, but decays in about a week when palbociclib is removed, so that the growth during a palbociclib interval of the alternating treatment is similar to its growth during the first week of palbociclib monotreatment (Figures S8A and S8B). While the difference in proliferation between mono and alternating treatments was not dramatic during the first two weeks, it became significant thereafter, with the alternating treatment cells approximately doubling every week (average 33.8-fold increase at week 5, 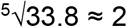, average 52.3-fold increase during the second 5 weeks, 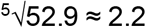). Part of the reason for this is that the palbociclib intervals of the alternating therapy do not experience the slowdown of the constant palbociclib cells to the same extent. But the other reason is that the growth during the ICI intervals is much greater than that of ICI monotherapy. To look for a mechanistic reason for the excessive proliferation, we measured the protein levels of cyclinD1, Cdk4, Cdk6, cyclinE1 and Cdk2 on days 35 and 70 for each arm of the experiment. Palbociclib treatment in both arms significantly increased the expression of cyclinD1 (Figure 4E). The increase of cyclinD1 during palbociclib treatment may be due to different degradation rates between the cyclinD1:Cdk4/6: palbociclib complex (number 14 in Table 1) and the cyclinD1:Cdk4/6 (number 12 in Table 1). When calibrating the model, we allowed the degradation rate of cyclinD1:Cdk4/6:palbociclib to be smaller than the degradation rate of cyclinD1:Cdk4/6 but greater than the degradation rate of cyclinD1:Cdk4/6:p21. This results in accumulation of cyclinD1 during palbociclib treatment that can partly explain the increase of cyclinD1 in the constant palbociclib cases at 35d and 70d, as well as the alternating case at 35d.

**Table 1.** Model Variables and Initial Values. Variable names are italicized.

| Variable name | Description | Initial value | Half-life |
| --- | --- | --- | --- |
| (1) *E2<sub>media</sub>* | E2 concentration in the media | 10nM | - |
| (2) *E2<sub>cell</sub>* | E2 concentration in the cell | 10nM | - |
| (3) *ER* | Estrogen receptor $\alpha$ | 1.97nM | ~4-5h Wijayaratne et al., 2019. |
| (4) *E2ER* | Estrogen bound estrogen receptor $\alpha$ | 835.19nM | ~3-4h Wijayaratne et al., 2019 |
| (5) *E2NSB* | Estrogen bound non-specific binding | 6697.83nM | - |
| (6) *ICIER* | ICI 182,780 bound estrogen receptor | 0nM | < 3-4h Wijayaratne et al., 2019 |
| (7) *rescyclinD1palbo* | Variable induced by palbociclib increasing cyclinD1 | 0nM | - |
| (8) *cyclinD1* | Protein cyclinD1 | $0.62 \times 10^{-6}$nM | ~0.4h Alao 2007 |
| (9) *cdk46* | Protein Cdk4/6 | 3365.58 | ~5h Gabrielli et al., 1999 |
| (10) *cdk46palbo* | Palbociclib bound Cdk4/6 | 0nM | - |
| (11) *cdk46abema* | Abemaciclib bound Cdk4/6 | 0nM | - |
| (12) *cyclinD1cdk46* | CyclinD1 bound Cdk4/6 | 33.83nM | - |
| (13) *cyclinD1cdk46p21* | p21 bound cyclinD1:Cdk4/6 | 21.84nM | - |
| (14) *cyclinD1cdk46palbo* | Palbociclib bound cyclinD1:Cdk4/6 | 0nM | - |
| (15) *cyclinD1cdk46abema* | Abemaciclib bound cyclinD1:Cdk4/6 | 0nM | - |
| (16) *cyclinD1cdk46p21palbo* | Palbociclib bound cyclinD1:Cdk46:p21 | 0nM | - |
| (17) *cyclinD1cdk46p21abema* | Abemaciclib bound cyclinD1:Cdk46:p21 | 0nM | - |
| (18) *cMyc* | Protein c-Myc | 9.75nM | ~0.3h Gregory et al., 2000 |
| (19) *p21* | Protein p21 | 0.0027nM | ~0.3-1h Abbas et al., 2009 |
| (20) *cyclinE* | Protein cyclinE | 0.16nM | ~0.5h Singer et al., 1993 |
| (21) *cyclinEp21* | p21 bound cyclinE | 0.036nM | - |
| (22) *Rb* | Retinoblastoma protein | 53.01nM | ~2-3h Oh et al., 2010 |
| (23) *pRb* | Hypophosphorylated RB1 (RB1-p) | 16.64nM | ~2-3h Oh et al., 2010 |
| (24) *ppRb* | Hyperphosphorylated RB1 (RB1-pp) | 0.49nM | > 4h Oh et al., 2010 |
| (25) *respropalbo* | Variable induced by palbociclib inhibiting proliferation | 0nM | - |
| (26) *Nalive* | Alive cell number | 0.93<sup>a</sup> | - |
| (27) *Ndead* | Dead cell number | 0.07<sup>a</sup> | - |

<sup>a</sup>The values of *Nalive* and *Ndead* are assigned according to the alive and apoptotic percentage in Figure S5.

However, we also saw that cyclinD1 experimentally increases at 70 days in the alternating treatment, which is just finishing an ICI interval. This increase cannot be explained by the decreased degradation rate of cyclinD1:Cdk4/6:palbociclib as this effect rapidly decays during the ICI interval (Figure S8E). In order to account for this effect, we added a phenomenological variable, *rescyclinD*1*palbo* (number 7 in Table 1), to the model that gradually increased cyclinD1 in response to long-term palbociclib treatment. The effect decreases slowly once palbociclib is removed (Figure S8C and S8D), so that the cyclinD1 levels during the ICI intervals of the alternating treatment are increased over the levels in the ICI monotreatment (Figure 4B and 4E). The above changes to the model enabled it to capture the proliferation under alternating therapy as shown in Figure. 4A. The cyclinD1 level is higher during the ICI intervals of the alternating treatment compared to ICI monotreatment (Figure 4B), which results in higher cyclinD1:Cdk4/6 and RB1-pp levels (Figure 4C and 4D). This effect causes the growth after the cells are transitioned from palbociclib to ICI to be greater than would otherwise be expected. The rapidly decaying peaks of cyclinD1:Cdk4/6 and RB1-pp seen at the palbociclib to ICI transition are due to the sudden release of palbociclib free Cdk4/6 and its complexes after palbociclib withdrawal.

### Protein Changes at 10 Weeks

Increased levels of the five proteins we measured, Figures 4E–4I are all associated with palbociclib resistance in the literature (Herrera-Abreu et al., 2016; Portman et al., 2018; Hafner et al., 2019; Knudsen et al., 2020; Pandey et al., 2020). In our experiment, Cdk6, cyclinE and Cdk2 levels show no statistically significant difference among the different treatment conditions. Although Cdk4 does show a statistically significant increase compared to untreated cells, the up-regulation is small (mean value of 1.4 at alternating treatment 35 days and 1.3 at palbociclib monotreatment at 70 days). Only cyclinD1 shows a large increase compared to untreated cells.

There is no significant difference in cyclinD1 level between palbociclib monotreatment and the alternating treatment during palbociclib intervals in Figure 4E. Moreover, in order to test whether the cyclinD1 gradually increases in response to long-term palbociclib treatment, as would be expected of a long term resistance mechanism, we measured cyclinD1 changes at 7 days and 14 days after 750nM palbociclib treatment. Figure 4J shows that the cyclinD1 level is already upregulated at 7 days and there are no significant differences in cyclinD1 levels among palbociclib monotreatment at 7 days, 14 days, 35 days, or 70 days. The observed increases in cyclinD1 can be explained by a rapid response to palbociclib treatment and do not represent a long-term change leading to resistance. Therefore, the five quantified proteins do not indicate any difference in moving towards resistance to palbociclib between the mono and alternating treatments.

### Palbociclib Dose-Response Changes at 10 Weeks versus 12 months

At the end of 10 weeks, a 7-day palbociclib dose-response assay was used to compare the proliferation of MCF7 cells after undergoing no treatment, monotreatment or alternating treatment. Figures 5A–5C show the results for three different normalizations: growth in vehicle, number of initial cells at t=0, and the growth rate inhibition metric, GR (Hafner et al., 2016). Figure 5A normalizes the proliferation of each case to its proliferation in vehicle, which is the usual method of normalization in biological experiments. From this plot we see that the alternating treatment cells are much more sensitive to palbociclib, compared with the monotreatment cells, at all doses of palbociclib. This would lead one to think that the alternating treatment is producing less resistant cells compared to monotreatment. However, when the dose-response results are normalized to t = 0, as shown in Figure 5B, we see that the proliferation of the palbociclib monotreatment cells is much less than that of the vehicle treated (E2 control). Because the proliferation is already low, and palbociclib does not significantly upregulate apoptosis (Roskoski 2016), the proliferation cannot decrease much further, making the cells appear less sensitive to palbociclib when normalized to vehicle. In contrast, the cells from alternating treatment have a relatively higher proliferation in vehicle and palbociclib can inhibit the proliferation more, which makes the cells appear sensitive to palbociclib. It should be noted, however, that for all doses, the alternating treatment cells proliferate faster than the monotreatment cells, which makes it impossible to claim an advantage for alternating treatment at 10 weeks, even if by standard measures the alternation results in cells that are more sensitive to palbociclib.

**Figure 5.**
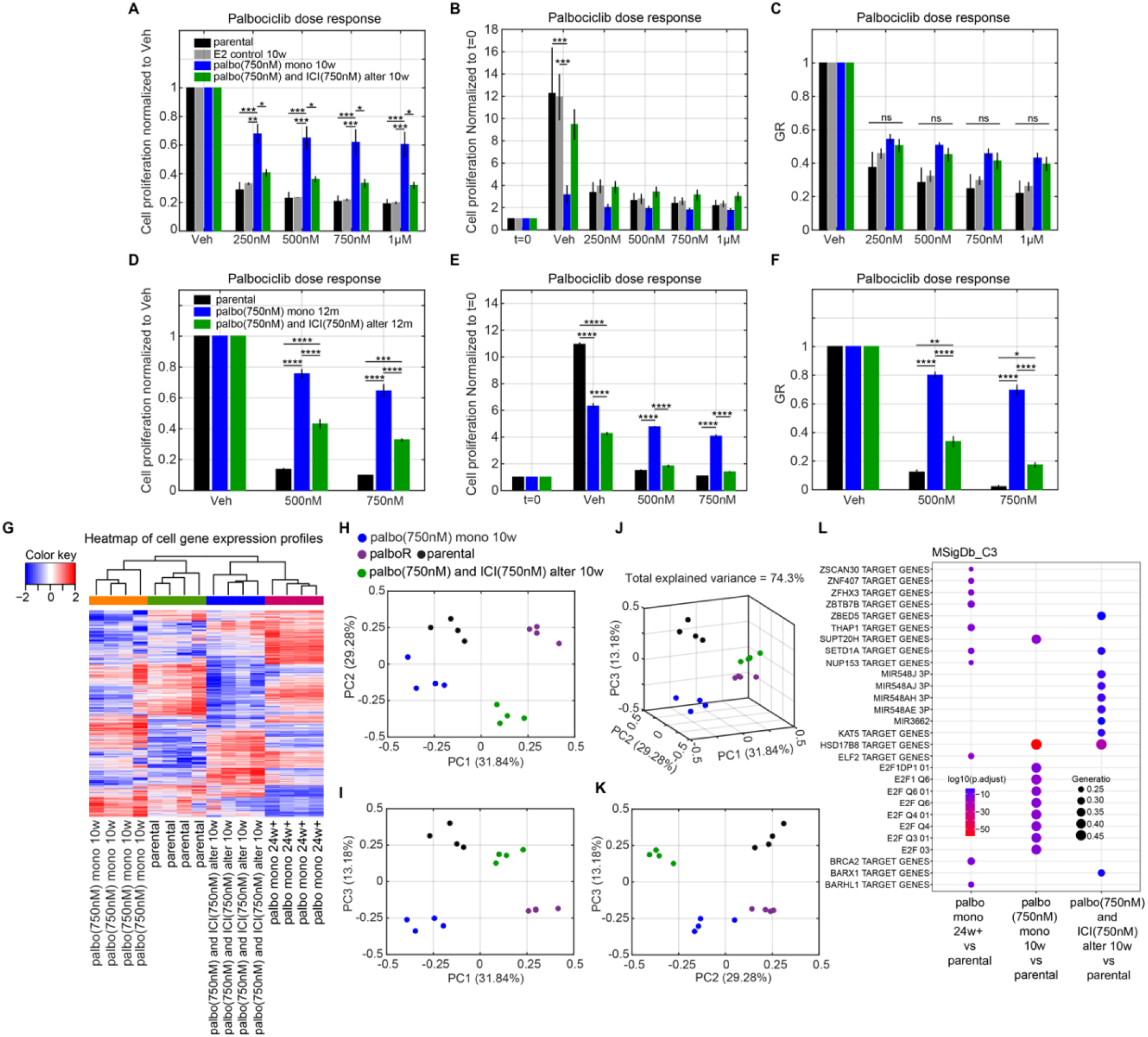
Palbociclib Dose Response and Gene Expression Profiles for Cells after Long Time Mono and Alternating Treatments. (A) Palbociclib dose response normalized to vehicle on cells after 10 weeks palbociclib (750nM) monotreatment and alternating treatment compared to parental MCF7 cells and MCF7 cells in 10 weeks E2 control condition. The alternating treatment is the same as Figure 4A, which is E2+palbo(750nM) alternating with E2+ICI(750nM). Each treatment period is 7days and starts with E2+palbo(750nM). The cells in all conditions are re-plated at 35days and the dose responses are tested at 70 days. (B) Palbociclib dose response normalized to t=0, otherwise same as (A). (C) The GR value of palbociclib dose response, otherwise same as (A). (D) Palbociclib dose response normalized to vehicle for cells after 12 months palbociclib (750nM) monotreatment and alternating treatment compared to parental MCF7 cells. Treatments are the same as (A) except the alternation period is 1 month, the duration is extended to 12 months, and the dose responses are tested at 12 months. (E) Palbociclib dose response normalized to t=0, otherwise same as (D). (F) The GR value of palbociclib dose response, otherwise same as (D). (G) Heatmap of gene expression profiles for cells after 10 weeks palbociclib monotreatment, cells after 10 weeks alternating treatment, parental MCF7 cells and cells cultured over 6 months in palbociclib (500nM). The cells from palbociclib monotreatment and alternating treatment are the same as (A). (H) Principal component analysis of gene expression profiles on the same cells as (G). (PC1 vs PC2). (I) Principal component analysis of gene expression profile on the same cells as (G). (PC1 vs PC3). (J) Principal component analysis of gene expression profile on the same cells as (G). (PC2 vs PC3). (K) Principal component analysis of gene expression profile on the same cells as (G). (PC1 vs PC2 vs PC3). (L) Gene Set Enrichment Analysis (GSEA) was performed on the same cells as (G). The C3 regulatory target gene sets in the Molecular Signatures Database (MSigDB) were used.

This problem of interpretation has been noticed previously and drove the development of a new metric, growth-rate inhibition (GR, see STAR Methods). If control cells undergo different numbers of divisions because of a difference in proliferation rate, the dose-response values will vary dramatically (Hafner et al., 2016). The standard dose response values are sensitive to the basal proliferation rate and GR accounts for this by normalizing the proliferation rate under treatment to the proliferation rate of the control (Hafner et al., 2016). GR is robust to variations in cell growth rate and quantifies the efficacy of a drug on a per-division basis, which can ensure that fast- and slow-dividing cells responding equally to a drug are scored equivalently (Hafner et al., 2016). Figure 5C shows the GR values for the palbociclib dose response and there is no significant difference between the mono and alternating treatments at 10 weeks. Therefore, although the dose response normalized to vehicle, Figure 5A, shows a difference between mono and alternating treatment, this effect comes from the different basal cell division rates of the mono and alternating treatment and obscures the true nature of the palbociclib dose response.

To explore what happens at a longer time scale, a 12 month alternating experiment using the same drugs and doses was performed. The alternation took place at the end of each month when the cells were also replated. At the end of 12 months a dose response was performed and the results are shown in Figures. 5D–5F. At this time, the palbociclib monotreatment cells were outgrowing the alternating cells in vehicle (Figure. 5E), but the growth of each arm was more similar than at 10 weeks. The result is that all three normalizations show similar behavior: the alternating cells are significantly more sensitive to palbociclib than the palbociclib monotreatment cells, indicating a delay in acquiring resistance. The alternating cells are beginning to acquire resistance, however, as can be seen by comparison to the parental cells in Figure. 5F. So, alternating therapies do show promise for delaying resistance, but better protocols are needed to hold down the excessive growth seen in the 10 week experiment.

### Gene Expression Changes at 10 weeks

Lastly, we analyzed gene expression profiles to look for differences between the palbociclib mono and alternating treatment cells at 10 weeks. Figure 5G shows the heatmap of differentially expressed genes for four cases of MCF7 cells: parental cells (control), 10 weeks of palbociclib monotreatment, 10 weeks of alternating treatment and cells cultured for >6 months (palbo mono 24w+) in palbociclib (500 nM). Although the alternating treatment cells clustered with the palbo mono 24w+ cells, the heatmap revealed distinct expression patterns for the four different treatments. The reason that alternating cells are in the same cluster with the palbo mono 24w+ cells is likely because they both have positive values of the first principal component (PC1), as shown by principal component analysis in Figures 5H–5K. The 2D and 3D principal component plots clearly show that cells under the four different treatments are separated into different groups. Gene Set Enrichment Analysis (GSEA) of the C3 regulatory target gene sets in the Molecular Signatures Database (MSigDB) is shown in Figure 5L. The first 10 most significantly different regulatory target gene sets are plotted. Under the alternating treatment, the most changed gene sets are microRNA regulated, which might be caused by prolonged ICI treatment (Rao et al., 2011; Zhou et al, 2018; Guo et al., 2019). Under the palbociclib monotreatment, the E2F regulated gene sets are the most changed. The E2F transcription factor is the central player in regulating the expression of genes involved in the G1 to S phase transition and the target genes in the listed sets include cyclinD1, cyclinE, Cdk2, Cdc25A, cyclinA, etc. (Ren et al., 2002; Stevens et al., 2003). In the palbo mono 24w+ cells, different gene sets are altered compared to the 10 week mono and alternating treatment cells, which might be related to the ongoing development of resistance such as BARHL1 target genes (Lim et al., 2022).

### Using Model-Generated Isobolograms to Determine Synergies

Cancer cells depend on a variety of molecular mechanisms for proliferation or survival, and therefore, drug combinations are often used to simultaneously target key molecular mechanisms to more effectively reduce proliferation, or help delay or overcome resistance (Narayan et al., 2020). A key question for drug combinations is whether there is a synergism between the drugs. A synergistic interaction between drugs may allow significantly lower doses of the individual drugs when used in combination as opposed to individually. It may benefit patients by reducing toxicity and adverse effects. There are numerous ways to define drug synergy, but we make use of the isobologram as we think it gives the clearest picture of the interaction of two drugs. An isobologram is a graph of lines of constant effect, called isoboles, proposed by Loewe in 1953 (Tallarida 2011). The isobologram is based on drug effect measurements made for a large number of different doses for the drugs, given both individually and in combination.

The upper plot of Figure 6A illustrates an ideal sampling scheme, where each axis represents the dose for a specified drug. Each blue hexagon is a measurement of the effect either solely from drug 1, or solely from drug 2, or of the combination effect from the doses of drug 1 and drug 2 that make up its coordinates. By drug effect, we mean the value of some measurable attribute of interest. It might be the percent of apoptotic cells in a culture, the cellular proliferation rate, the reduction in tumor size, the toxicity, or some other physiological index such as heart rate or blood pressure. Any measurable metric of interest can be used to defined an isobologram. After measuring the effect at each dosage point of the isobologram, we draw the isoboles, which are lines joining the points of equal measured effect. The lower plot of Figure 6A shows example isoboles, where the different drug doses at each point on the isoboles give the same effect. The isobologram simply reduces the 3-D plot of drug effect versus drug 1 and drug 2 to a 2-D contour plot that is easier to interpret quantitatively. The relationship between the two drugs can be defined from the shape of the isoboles, as illustrated in lower plot Figure 6A:

**Figure 6.**
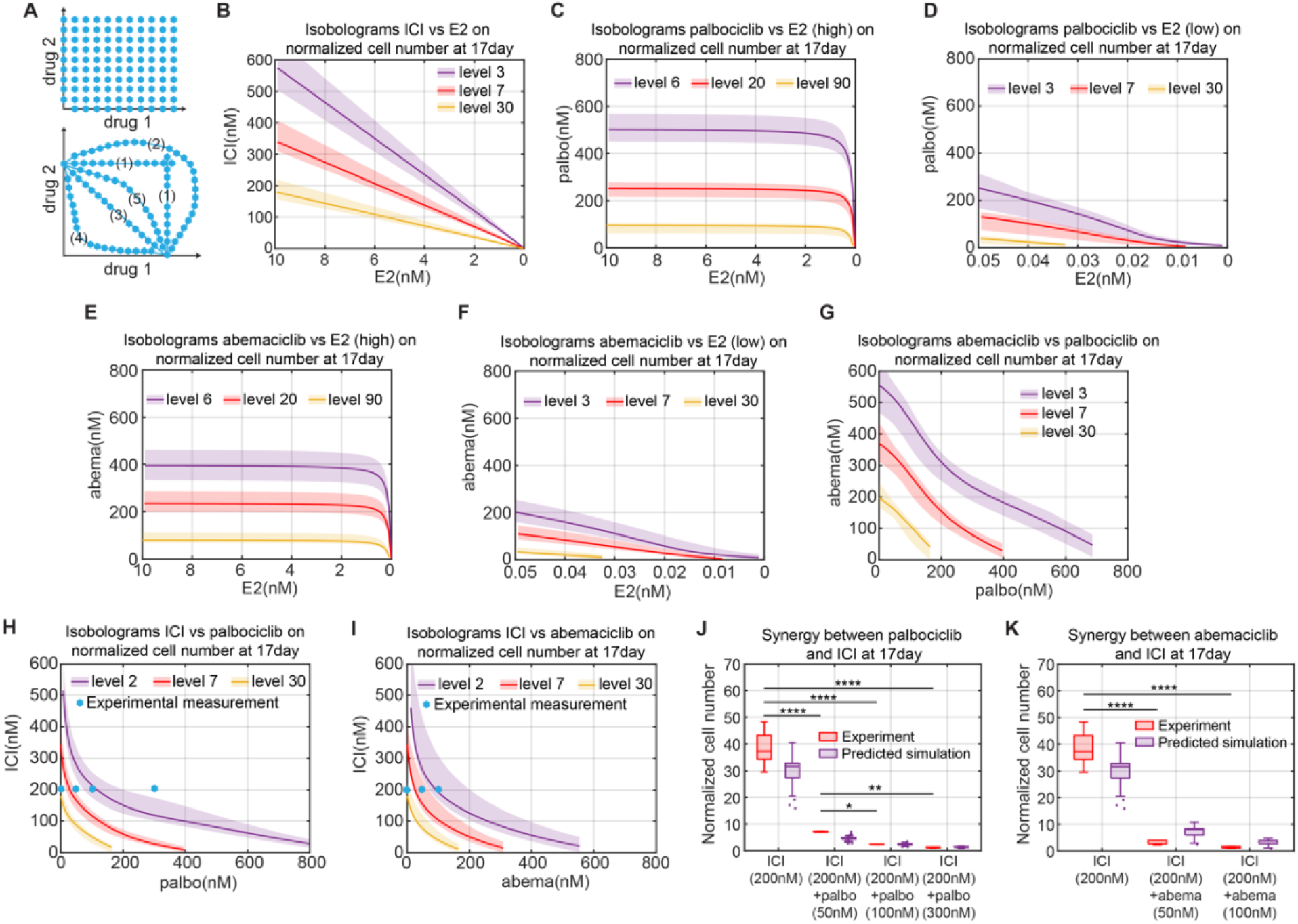
Model Simulation of Isobologram among Various Treatment Methods and Experimental Verifications. (A) Illustration of the isobologram. Each blue hexagon represents a measurement point for mono or combination drug treatment effects. The lines joining the (interpolated) points of equal measured effect are isoboles, such as lines in the lower plot, which represent different interaction types: (1) Independence; (2) Antagonism; (3) Additive; (4) Super-additive; (5) Sub-additive. (B) Model simulation of isobologram between ICI and E2 for the normalized cell number at 17days. Different colors of the isobole represents the different levels of normalized cell number. The solid line represents the lowest cost value simulation and the shaded regions contain the central 98% of the cohort simulations. (C) Model simulation of isobologram between palbociclib and E2 (high concentration) for the normalized cell number at 17days. (D) Model simulation of isobologram between palbociclib and E2 (low concentration) for the normalized cell number at 17days. (E) Model simulation of isobologram between abemaciclib and E2 (high concentration) for the normalized cell number at 17days. (F) Model simulation of isobologram between abemaciclib and E2 (low concentration) for the normalized cell number at 17days. (G) Model simulation of isobologram between palbociclib and abemaciclib for the normalized cell number at 17days. (H) Model simulation of isobologram between palbociclib and ICI for the normalized cell number at 17days. (I) Model simulation of isobologram between abemaciclib and ICI for the normalized cell number at 17days. (J) Boxplot of the model predictions and experimental verifications of normalized cell number showing the synergism between palbociclib and ICI. The doses of drug combinations used in the experiment are marked by the blue hexagons in (H). The prediction results shown in purple are from all cohort simulation results. Statistical testing was performed by two-way ANOVA (ns: non-significant; *: p <0.05; **: p ≤ 0.01; ***: p ≤ 0.001; ****: p ≤ 0.0001). Center line on each box is the median. The bottom and top lines on each box are the 25^th^ and 75^th^ percentiles, respectively. The whiskers are maximum and minimum values without considering outliers. Data points are considered outliers if they are more than 1.5×IQR (interquartile range) below the 25^th^ percentile or above the 75^th^ percentile. (K) Boxplot of the model predictions and experimental verifications of normalized cell number showing the synergism between abemaciclib and ICI. The doses of drug combinations used in the experiment are marked by the blue hexagons in (I). The prediction results shown in purple are from all cohort simulation results. The statistical testing used and explanation of the boxplot are the same as (J).

#### Independence

Isoboles (1) show an independent relationship between drug 1 and drug 2. If drug 2 has no effect the isoboles are vertical, and if drug 1 has no effect they are horizontal.

#### Antagonism

Isobole (2) illustrates an antagonistic relationship between drug 1 and drug 2. Combining drug 2 with drug 1 requires increasing the dose of drug 1 to have the same effect and vice versa.

#### Additive

Isobole (3) shows an additive relationship between drug 1 and drug 2, meaning that the combined effect of the two drugs is consistent with their individual potencies. Put another way, if d is the dose of a drug and D is the dose of that drug producing the specified effect as a monotreatment, then an additive effect means that a change in the normalized dose of drug 1, d1/D1, requires an equal and opposite change in the normalized dose of drug 2, d2/D2, in order to preserve the specified effect.

#### Super-additive

An isobole positioned below where an additive isobole would be, illustrated by isobole (4), is called superadditive (Huang et al., 2019).

#### Sub-additive

An isobole positioned above where an additive isobole would be, illustrated by the isobole (5), is called subadditive (Huang et al., 2019).

In this paper, we define a drug combination to be synergistic if it is super-additive. To obtain accurate isoboles, a large number of measurements are needed, as shown in Figure 6A. This makes the experimental determination of isoboles a challenging project. With a mathematical model, however, the generation of isoboles is essentially trivial, as a large number of simulations can easily be run and the results provided to a contour plotting program to get the isoboles. Figure 6B–6I shows the isoboles computed by our model for cases ICI v. E2, palbociclib v. E2 (high), palbociclib v. E2 (low), abemaciclib v. E2 (high), abemaciclib v. E2 (low), abemaciclib v. palbociclib, ICI v. palbociclib and ICI v. abemaciclib, respectively. The drug effect considered in these isoboles is the fold-change in cell number over 17 days of treatment and the results illustrate a range of different interaction types.

Figure 6B shows that the interaction between ICI and –E2 is additive. This is reasonable because both ICI and –E2 target the estrogen signaling pathway and decrease the E2:ER transcription factor level without directly influencing any other targets in the model. ICI and –E2 influence ER only through binding and unbinding reactions, so the level of E2:ER will linearly decrease after increasing ICI or decreasing E2 (Figure S7). Therefore, the effects of ICI and –E2 as mono and combination treatments are the same, to linearly decrease E2:ER level.

Figure 6C–6D shows the interaction between palbociclib and –E2 and indicates that the effect of palbociclib is largely independent of the concentration of E2 until the E2 concentration gets into the picomolar range. Figure 6D provides a zoomed in plot of the isoboles for low concentrations of E2 and shows that the interaction between palbociclib and deprivation is additive or slightly super-additive in this region.

Figure 6E–6F shows the interaction between abemaciclib and –E2. As expected, the interaction between abemaciclib and –E2 is same as palbociclib with –E2, which is independent of the concentration of E2 until the E2 concentration gets into the picomolar range, where the interaction becomes additive or slightly super-additive. Figure 6G shows the interaction between abemaciclib and palbociclib is primarily additive. This is reasonable because abemaciclib and palbociclib both target the Cdk4/6 activity with a binding-unbinding reaction.

Figure 6H shows the interaction between ICI and palbociclib and indicates a significant synergism between ICI and palbociclib. To test this synergy predicted by the model, an experiment was performed where the ICI dose was held constant at 200nM and various doses of palbociclib were added (0nM, 50nM, 100nM and 300nM, blue hexagon in Figure 6H). The isoboles in Figure 6H predict that we will see a dramatic decrease in population growth, which is borne out in the experimental results shown in Figure 6J. It should be emphasized that the model parameters were calibrated using only data from ICI and palbociclib monotreatments, not data from combination treatments. We believe the reason the model gives an experimentally consistent prediction of this significant synergism is because the structure of the model is based on the dominant signaling pathways of the system. In our mechanistic model, we include ICI’s effects on E2:ER, E2:ER’s effects on cyclinD1, and palbociclib’s effects on Cdk4/6. Therefore, the activity of the cyclinD1:Cdk4/6 kinase is attacked from both the cyclinD1 and Cdk4/6 directions to create the synergism. This may be the reason that palbociclib in combination with endocrine therapies achieved substantial improvement in survival outcomes in clinical trials and quickly became the first-line choice of treatment for ER+ breast cancer (Xi et al., 2020). This synergy is in contrast to the combination of ICI and –E2, whose mechanisms both target E2:ER, and produce an additive but not synergistic response. Figure 6I shows the interaction between ICI and abemaciclib which also indicates a significant synergism. Likewise, experiments were performed to test this synergy predicted by the model where the ICI dose was held constant at 200nM and various doses of abemaciclib were added (0nM, 50nM, 100nM, blue hexagons in Figure 6I). As expected, the synergistic response predicted in Figure 6I is borne out in the experimental results shown in Figure 6K. The explanation for the synergism between abemaciclib and ICI is same as for palbociclib and ICI.

The ability to easily produce isoboles for various metrics, such as proliferation over a specified time frame, allows us to propose optimal combination therapies. For example, considering a combination treatment of ICI and palbociclib, we can minimize the total dose of drugs, [ICI]+[palbociclib], that achieves our specified objective. Other possibilities include minimizing the total normalized dose of the drugs or some other weighted dose, [ICI]+λ[palbociclib], that reflects preferences based on toxicity or other concerns. Since palbociclib is typically used in the clinic in an intermittent fashion, three weeks on and one week off, due to neutropenia concerns, we could limit the above optimizations to lower doses of palbociclib that allow constant application so that excessive proliferation during the week off is avoided.

### Alternating Treatment Predictions

Ultimately, as mentioned above, we would like to show that the mathematical model allows us to propose optimal combination therapies. The experimental proliferation results in Figure 4A show that alternating palbociclib with ICI produces dramatically greater proliferation than the monotreatment. So, even if this alternation results in cells that are less resistant, it would not be a viable therapeutic approach. On the other hand, continuously applied monotreatment almost always leads to resistance and recurrence. So we used the model to look for better options to the simplistic alternating treatment we used above.

Simply trying to minimize proliferation will lead to the unhelpful answer of massive drug doses that would never be tolerable in any real application. Therefore, we decided to minimize the total drug dose over a 12 week time period, subject to the constraint that the overall fold-change be no greater than that of palbociclib monotreatment. Since this would likely lead to simply applying the best combination of palbociclib and ICI continuously, leading ultimately to resistance, we specified the repeating cycle to consist of 1 week of palbociclib, 1 week of a combination, 1 week of ICI, and 1 week of the same combination again. An optimization routine choses the drug doses in each week so as to minimize the total drug concentration applied over the 12 week period. The results are shown in Figure 7A. By design, the alternating treatment has the same fold-change as the monotreatment, but the optimized alternating treatment uses about 1200nM less total drug dose per cycle compared to the palbociclib monotreatment, 1905nM to 3080nM, and about 900nM less total drug dose per cycle than the ICI monotreatment, 1905nM to 2800nM. The combination treatment intervals not only find the synergistic sweet spot, noted above, to virtually stop growth and allow basal apoptosis to reduce the population, but they reduce the proliferation during the monotreatment intervals compared to switching directly from one monotreatment to another (Figures 7B–7D). This result shows that more sophisticated alternating treatments may provide benefits in terms of reduced drug dose while not continuously applying the same regimen, possibly delaying the onset of resistance.

**Figure 7.**
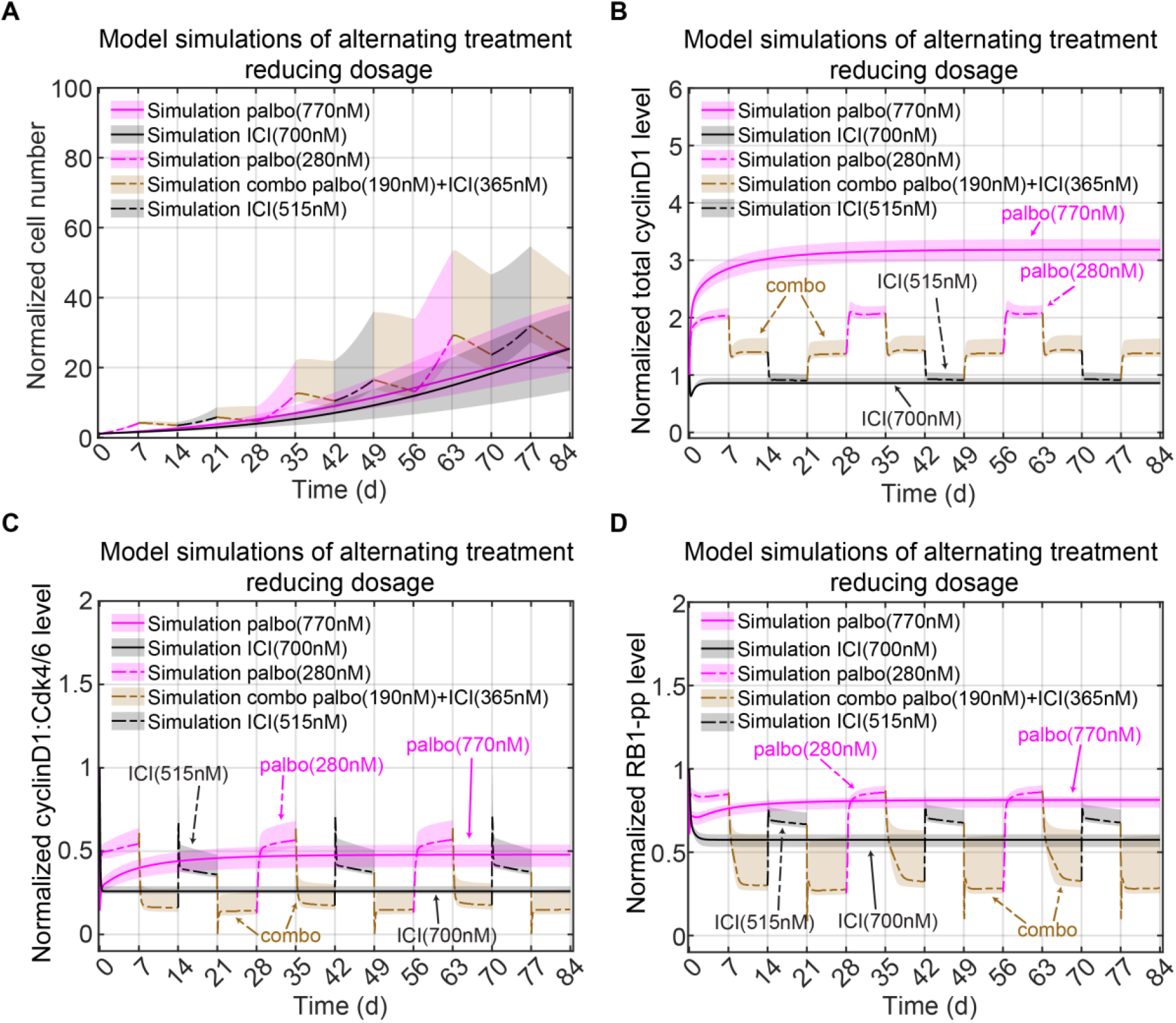
Optimal Treatment Design Using the Model. (A) Proposed Alternating treatment to reduce total drug dosage. E2+palbo(770nM) monotreatment is shown in purple with a solid line. E2+ICI(700nM) monotreatment is shown in black with a solid line. For the alternating treatment, each treatment period is 7days. In a 28 day cycle, the alternation starts with E2+palbo(280nM) shown in purple with a dashed line, then changes to a combination treatment of E2+palbo(190nM)+ICI(365nM) shown in a brown dashed line, then changes to E2+ICI(515nM) shown in a black dashed line, then changes to the combination treatment again. The cycle is repeated 3 times for a total of 84 days. The solid and dashed lines represent the lowest cost value simulation and the shaded regions contain the central 98% of the cohort simulations. (B) Model simulation of normalized total cyclinD1 level changes in the proposed alternating treatments shown in (A). The lines and shaded regions have the same meaning as (A). (C) Model simulation of normalized cyclinD1:Cdk4/6 level changes in the proposed alternating treatment shown in (A). (D) Model simulation of normalized RB1-pp level changes in the proposed alternating treatment shown in (A).

## Discussion

In this work, we built a mechanistic ODE model to capture the response of MCF7 cells to clinically used anticancer therapies for ER+ breast cancer. We used the model to recapitulate and predict drug treatment effects on these cells and optimize drug combinations. As the model has a mechanistic basis and the relevant targets were already included when creating the model for palbociclib, we showed that the model can be easily extended to test the effect of one of the other Cdk4/6 inhibitors, abemaciclib. We also illustrated the usefulness of the model to efficiently investigate synergism among the different treatments included in the model.

While much of the work in cell lines to explore the impact of therapies takes place over short time frames of less than a week, most clinical therapy occurs over much longer time frames of months and years (Serra et al., 2019; Gharib et al., 2022). The work reported here looks for insights from cell lines over these longer time periods. Because of the limited number of such experiments that can be run, trial and error approaches are not viable. We used a mathematical model of the system, calibrated on limited data, to guide our explorations and search for better therapy options. Predicting drug responses over long time periods is not simply a matter of taking a model calibrated on data from a week long experiment and running it for a longer period, as there are significant factors affecting the model that are only clearly seen over longer time periods. This necessitates long-term experiments to calibrate the model. One example of this is that cell proliferation under –E2 treatment over a long time scale behaved qualitatively different than proliferation over a short time scale. Another example of this is the excessive growth observed when ICI treatment was applied after initially treating with palbociclib. This observation and additional experiments led us to the fact that treatment with palbociclib increases cyclinD1, something we had missed earlier. The revised mathematical model allowed us to propose a protocol to counter this effect.

Since our ultimate goal is to delay or prevent the onset of drug resistance, adding resistance mechanisms to the model is a critical requirement for future work (Wander et al., 2020; Asghar et al., 2022; Pandey et al., 2022, Papadimitriou et al., 2022). The cyclinD1 change mentioned above is a minor step in that direction, but the development of resistance is a complex, multi-faceted process and there are many different pathways that lead to a drug resistant state (Lloyd et al., 2022; Watt et al., 2022). To see whether a therapeutic protocol delays the emergence of resistance compared to monotreatment will require experiments over time periods of many months, necessitating the use of mathematical models to propose the most promising protocols to explore (Serra etl al., 2019; Waller et al., 2019; Llombart-Cussac et al., 2021).

The use of alternating therapies to delay resistance is predicated on the assumption that the initial stages of acquiring resistance are reversible, which appears likely in many cases (Azizian et al., 2010; Cornell et al., 2019; Formisano et al., 2019; O’Brien et.al., 2020; De Angelis et al., 2021; Scheidemann et al., 2021; Sobhani et al., 2021; Kharenko et al., 2022). A critical mutation, however, can render most targeted therapies useless and thus upend any alternating protocol (O’Leary et al., 2018; Dustin et al., 2019; Nayar et al., 2019; Brett et al., 2021; O’Leary et al., 2021; Ono et al., 2021; Raimondi et al., 2021). To limit the probability of mutation, a much greater reduction in proliferation than is achieved in our current experiments is necessary. Periodically adding a more cytotoxic drug into the protocol is probably required (Das Thakur et al., 2013; Labrie et al., 2022). In addition, although alternating treatment does not continuously apply a single drug to attack the cancer cells, our current approach using standard of care treatments for ER+ breast cancer is to continuously arrest cells in the G1/S phase of the cell cycle with antiestrogens and Cdk4/6 inhibitors (Pernas et al., 2018). Resistant cells can bypass the G1/S blockade and alter G2/M cell cycle proteins to survive (O’Leary et al., 2018; Pancholi et al., 2020; Fallah et al., 2021; Ono et al., 2021). Therefore, targeting of multiple cell cycle phases may be needed to avoid development of resistance to current therapies in ER+ breast cancer (Aarts et al., 2012; Kettner et al., 2019; Portman et al., 2020; Pandey et al., 2022). Finally, we recognize that work in cell lines may not directly translate to animals and humans, but hope that it may provide insights that can benefit work closer to the clinic.

## METHODS

## ACKNOWLEDGMENTS

This work was partly supported by Public Health Service grant R01-CA201092 to W.T.B. and A.N.S.-H. Technical services were provided by shared resources at Georgetown University Medical Center, including the Tissue Culture Core Shared Resources and the Genomics and Epigenomics Shared Resources, that were funded through Public Health Service award 1P30-CA-51008 (Lombardi Comprehensive Cancer Center Support Grant). We also thank the Georgetown Breast Cancer Advocates (GBCA) for a patient’s perspective for this study.

## AUTHOR CONTRIBUTIONS

Conceptualization, W.T.B., A.N.S.-H., W.H.; methodology, W.T.B., A.N.S.-H., W.H.; software, W.H.; investigation, W.T.B., W.H.; data curation, D.M.D., Y.F., I.S.C.; writing–original draft, W.H., W.T.B.; writing–review & editing, W.T.B., A.N.S.-H; visualization, W.H.; supervision, A.N.S.-H., W.T.B.; project administration, W.T.B.

## EXPERIMENTAL MODEL AND SUBJECT DETAILS

| REAGENT or RESOURCE | SOURCE | IDENTIFIER |
| --- | --- | --- |
| Antibodies |  |  |
| anti-β-Actin | Santa Cruz Biotechnology | #sc-47778 |
| anti-β-Tubulin I | Sigma-Aldrich | #T7816 |
| anti-Cdk2 | Cell Signaling Technology | #2546 |
| anti-Cdk4 | Cell Signaling Technology | #12790 |
| anti-Cdk6 | Cell Signaling Technology | #3136 |
| anti-CyclinD1 | Cell Signaling Technology | #2978 |
| anti-CyclinE (HE12) | Santa Cruz Biotechnology | #sc-247 |
| anti-c-Myc | Cell Signaling Technology | #5605 |
| anti-ERα | ThermoFisher Scientific | #MA5-14104 |
| anti-RB1 | Cell Signaling Technology | #9309 |
| anti-RB1-pp(S612) | Aviva Systems Biology | #OAAB16108 |
| Chemicals, peptides, and recombinant proteins |  |  |
| phenol red-free improved minimal essential medium | Life Technologies, Grand Island, NY | A10488-01 |
| Faslodex/Fulvestrant; ICI182,780 | Tocris Bioscience, Ellisville, MO | S1191 |
| palbociclib | Tocris Bioscience, Ellisville, MO | S1116 |
| abemaciclib | Cayman Chemical | #17740 |
| Bovine Calf Serum Charcoal Stripped (CCS) | GeminiBio | #100-213 |
| 17β-estradiol (E2) | Sigma-Aldrich, St. Louis, MO | #E8875 |
| Complete Mini protease inhibitor mixture tablets | Roche Applied Science | 11836153001 |
| Apotracker | Thermofisher Scientific<br>Waltham, MA | 427402 |
| propidium iodide | Thermofisher Scientific<br>Waltham, MA | P1304MP |
| Critical commercial assays |  |  |
| bicinchoninic acid assay | Pierce | 23227 |
| RNeasy kit | Qiagen, Valencia, CA |  |
| Affymetrix GeneChip HG-U95 arrays | Affymetrix, Santa Clara, CA |  |
| Z1 Single Coulter Counter | Beckman Coulter, Miami, FL |  |
| Experimental models: Cell lines |  |  |
| MCF7 | Lombardi Comprehensive<br>Cancer Center, Georgetown<br>University, Washington, DC |  |

### Cell Culture and Reagents

MCF7 cells were obtained from Tissue Culture Shared Resources at Lombardi Comprehensive Cancer Center, Georgetown University, Washington, DC. MCF7 cells were grown in phenol red-free improved minimal essential medium (Life Technologies, Grand Island, NY; A10488-01) with 10% charcoal-stripped calf serum (CCS) and supplemented with 10nM 17β-estradiol (E2). ICI (Faslodex/Fulvestrant; ICI182,780) and palbociclib were obtained from Tocris Bioscience (Ellisville, MO). MCF7 cells were authenticated by DNA fingerprinting and tested regularly for Mycoplasma infection. All other chemicals were purchased from Sigma-Aldrich (St. Louis, MO).

## METHOD DETAILS

### Cell Proliferation Assay

Cells were seeded at a density of 4–5 × 10^4^ cells/well in 60 mm plates and treated with indicated drugs at 24 h post plating. E2 deprivation was obtained by washing cells 24 h post-plating (t = 0) with phosphate-buffered saline (PBS) and adding complete medium without E2 for the indicated times. To measure cell number at specific time-points, cells were trypsinized, resuspended in PBS and counted using a Z1 Single Coulter Counter (Beckman Coulter, Miami, FL).

### Western Blot Analysis

For Western blot analysis, cells were lysed for 30 min on ice with lysis buffer (50 mM Tris-HCl, pH 7.5, containing 150 mM NaCl, 1 mM EDTA, 0.5% sodium deoxycholate, 1% IGEPAL CA-630, 0.1% sodium dodecyl sulfate (SDS), 1 mM Na3VO4, 44 μg ml–1 phenylmethylsulfonyl fluoride) supplemented with Complete Mini protease inhibitor mixture tablets (Roche Applied Science). Total protein was quantified using the bicinchoninic acid assay (Pierce). Whole-cell lysate (20 μg) was resolved by SDS–polyacrylaminde gel electrophoresis.

### Apoptosis Assay

2-5 × 10^5^ cells were plated in 6-well plates, were treated for 72 h, and stained with Apotracker green and propidium iodide, respectively (Thermofisher Scientific Waltham, MA) according to the manufacturer’s protocol and fluorescence was measures by the Flow Cytometry Shared Resource at Georgetown University Medical Center. Each experiment was repeated at least three times.

### Microarray

Microarray analysis was performed using four biological replicates using Affymetrix HG U133 Plus 2.0 microarray at our Genomics and Epigenomics Shared Resources. Briefly, total RNA was extracted using the RNeasy kit (Qiagen, Valencia, CA, USA). RNA labeling and hybridization were performed according to the Affymetrix protocol for one-cycle target labeling. For each experiment, fragmented cDNA was hybridized in triplicates to Affymetrix GeneChip HG-U95 arrays (Affymetrix, Santa Clara, CA, USA).

### Dynamics of E2 Deprivation

Removing E2 completely from cultured cells that have been growing in medium containing E2 cannot be accomplished by simply changing to a medium containing no E2. The E2 deprivation procedure is conducted by exchanging the E2 medium with 5% charcoal stripped calf serum (CCS) and phenol-red free media (Lewis et al., 2005). The E2 level in CCS is routinely measured to be less than 4 pM (Lewis et al., 2005), equating to 0.2 pM in 5% CCS media. But the E2 in the cell is at a significantly higher concentration than that in the medium and it can diffuse back into the medium and cause an increase in the E2 concentration. While the concentration of E2 might be low, its effect might not be negligible because a direct mitogenic effect of exogenous E2 on MCF7 can be initiated as low as 3 pM and maximized at 0.2 to 10nM (Furuya et al., 1989). Furthermore, other than estrogen receptors, there exist nonspecific bindings between estrogen and other elements inside the cell (Strobl et al., 1979). Therefore, MCF7 cells growing in an E2 condition have a much higher internal concentration of E2 than that of medium due to non-specific binding of E2 in the cytoplasm as well as specific binding of E2 to various estrogen receptors in the cell. When we deprive the medium of estrogen, E2 from the cells leaches into the new medium and a new balance between the estrogen levels inside and outside the cell is achieved. The newly established E2 level that can be significant for maintaining proliferation. From the –E2 proliferation result shown in Figure 1D, the initial growth period is short and the cells nearly stop growing later on. As the medium is replaced as the experiment proceeds, the E2 level continues to drop and the cells stop proliferation.

In a one-week proliferation experiment (Figure S3), we changed the medium at time zero and at day 3 to –E2 medium and counted the cells on day 7. In a parallel experiment, an extra medium change was inserted at 3 hours. The experiment was conducted with two different plating densities. We can see that the extra media change, which further decreases the residual E2 level, significantly reduces the overall MCF7 proliferation at 1 week. Not only do the changes in E2 concentration with each successive medium change impact proliferation for long time continuous –E2 treatment, but these changes are also critically important when we consider alternating treatments. For example, if we alternate E2+palbo with –E2 treatment, after the transition from E2 to –E2 medium excess E2 will leach into the medium causing undesired growth. This issue drove us to model the E2 concentration dynamically. Thus, modeling the E2 dynamics is needed to capture the effect of alternating treatment.

### Dynamic Modeling of E2 Deprivation

After a medium change, the total number (#) of E2 molecules should be constant, so the amount of E2 leaving the cell should be equal to the amount of E2 entering the medium and vice versa. The rate of change in the number of E2 molecules in the cell caused by diffusion is:

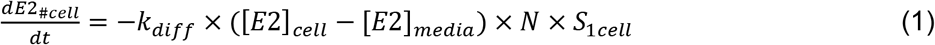

Where *E*2_#cell_ is the total number (#) of E2 molecules in the cells, *k_diff_* is the diffusion rate across the cell membrane and -*k_diff_* × ([*E*2]_*cell*_ – [*E*2]_*media*_) has units of #/(*m*^2^ × *t*), [*E*2]_*cell*_ is the E2 concentration in the cell, [*E*2]_*media*_ is the E2 concentration in the medium, *N* is the total cell number and *S_1cell_* is the surface area of a single cell.

Because *E*2_#*cell*_ = [*E*2]_*cell*_ × *Vol_cells_*, where *Vol_cells_* is the total volume of cells, which changes with time, equation (1) becomes:

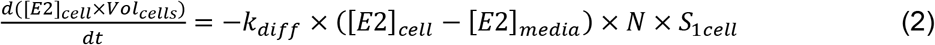

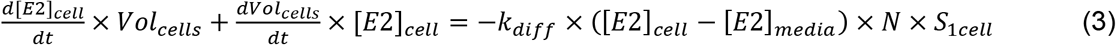

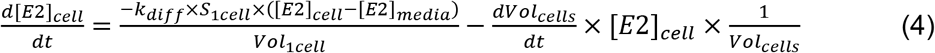

setting 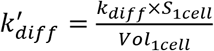, equation (7) becomes

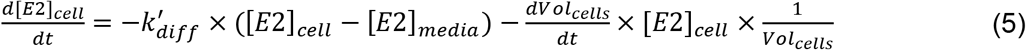

Where the first term on the right is the rate of change related to diffusion and the second term is the rate of change related to variations in total cell volume. To simplify the second term, note that:

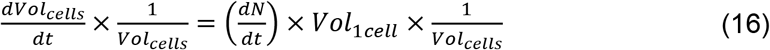

So equation (10) becomes

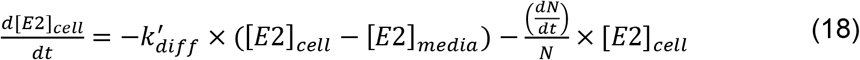

If we suppose the volume of the culture media doesn’t change, since it is massive compared to the total cell volume, then the rate of E2 concentration changes in the media becomes

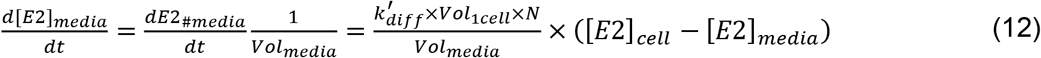

since the total number of molecules diffusing into the medium is equal to the total number of molecules diffusing out of the cells.

Then equation (18) and (12) are used to model the E2 dynamics during and after the deprivation. In Table 1, [*E*2]_*cell*_ is denoted as *E*2_*cell*_ and [*E*2]_*media*_ as *E*2_*media*_. Each time the medium is changed to –E2, the value [*E*2]_*media*_ is set to the value of *E*2_*dep*_ in Table 2. Each time the medium is changed to control condition, the value [*E*2]_*media*_ is set to the value of *E*2 in Table 2

**Table 2.**
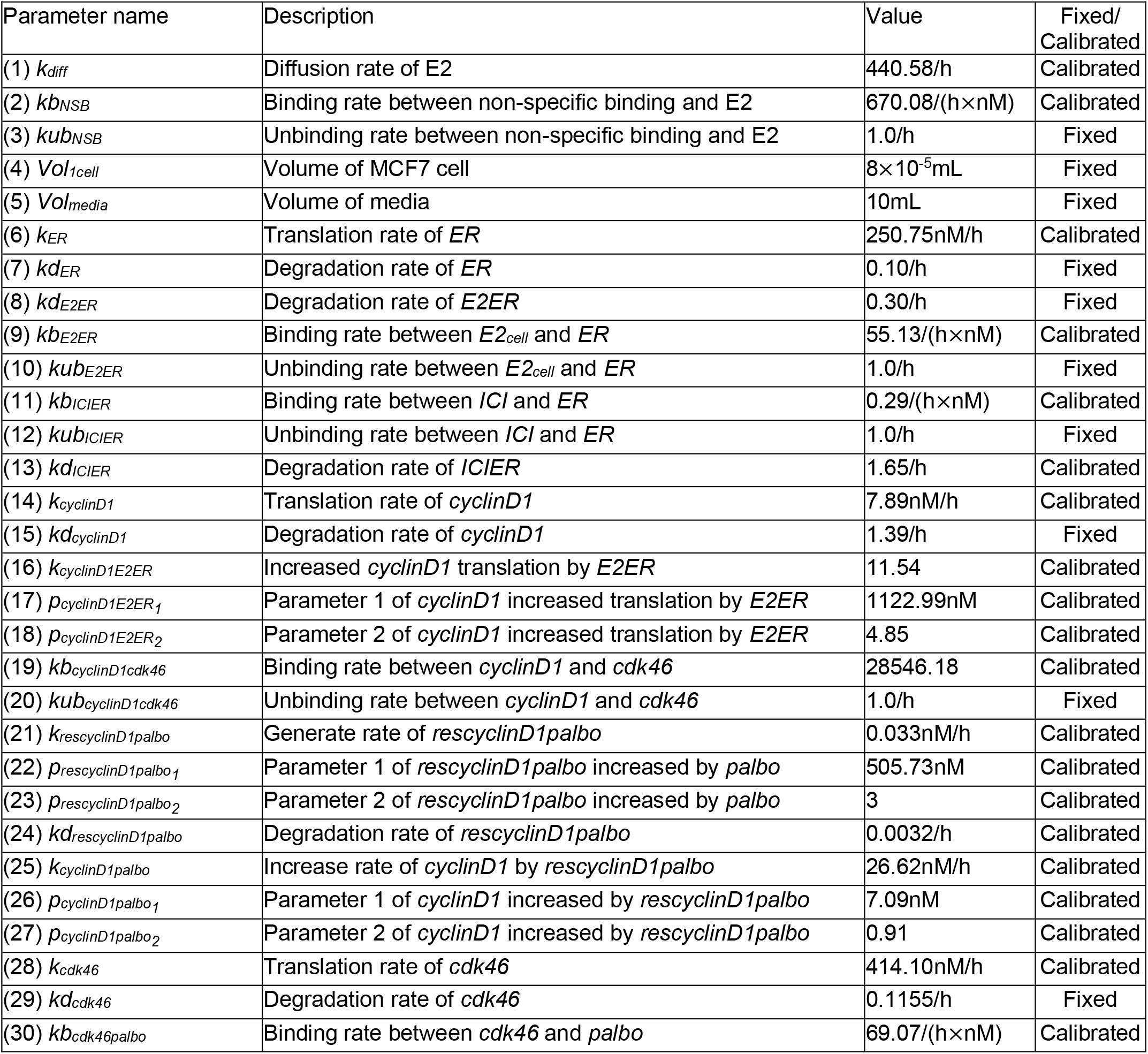
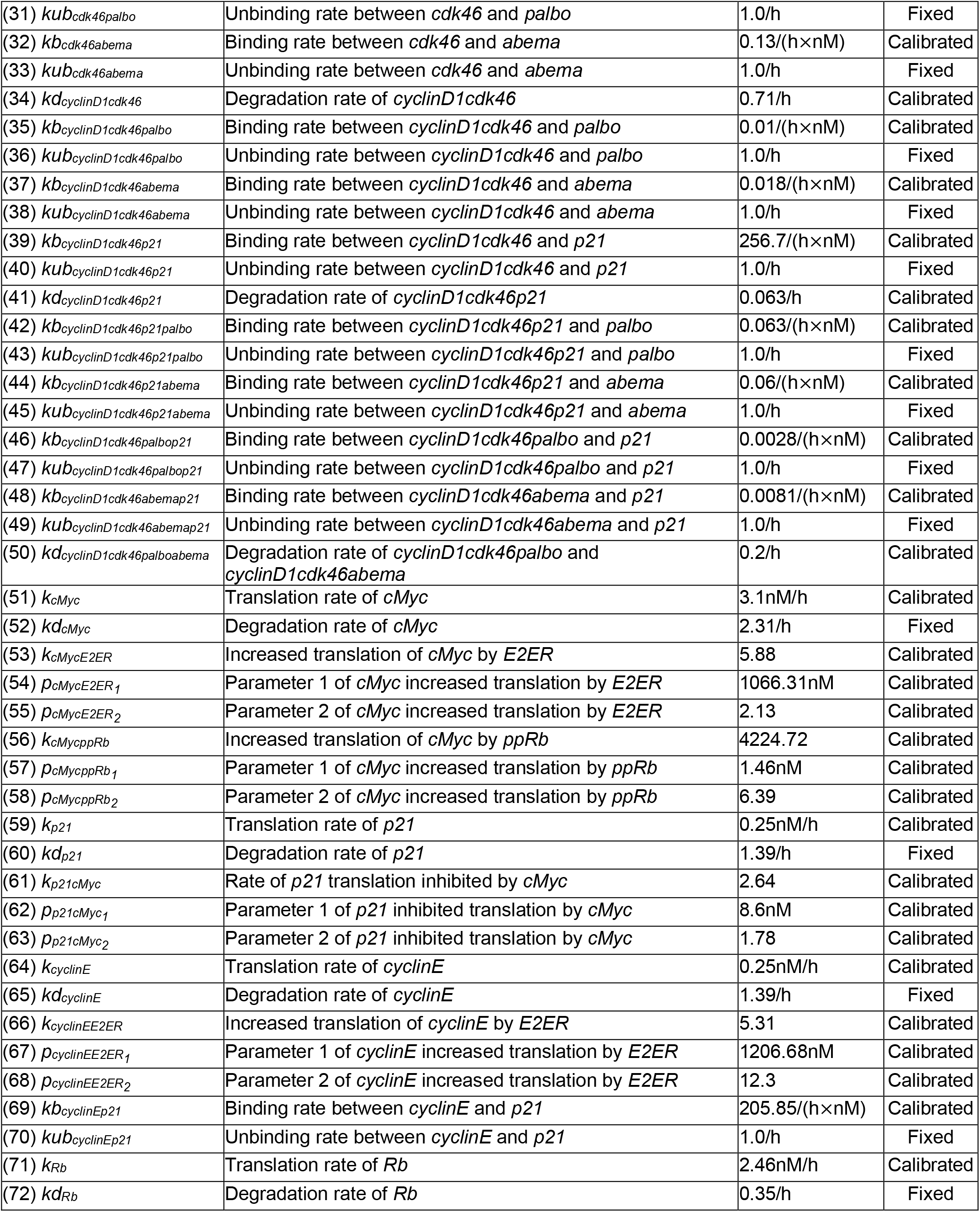
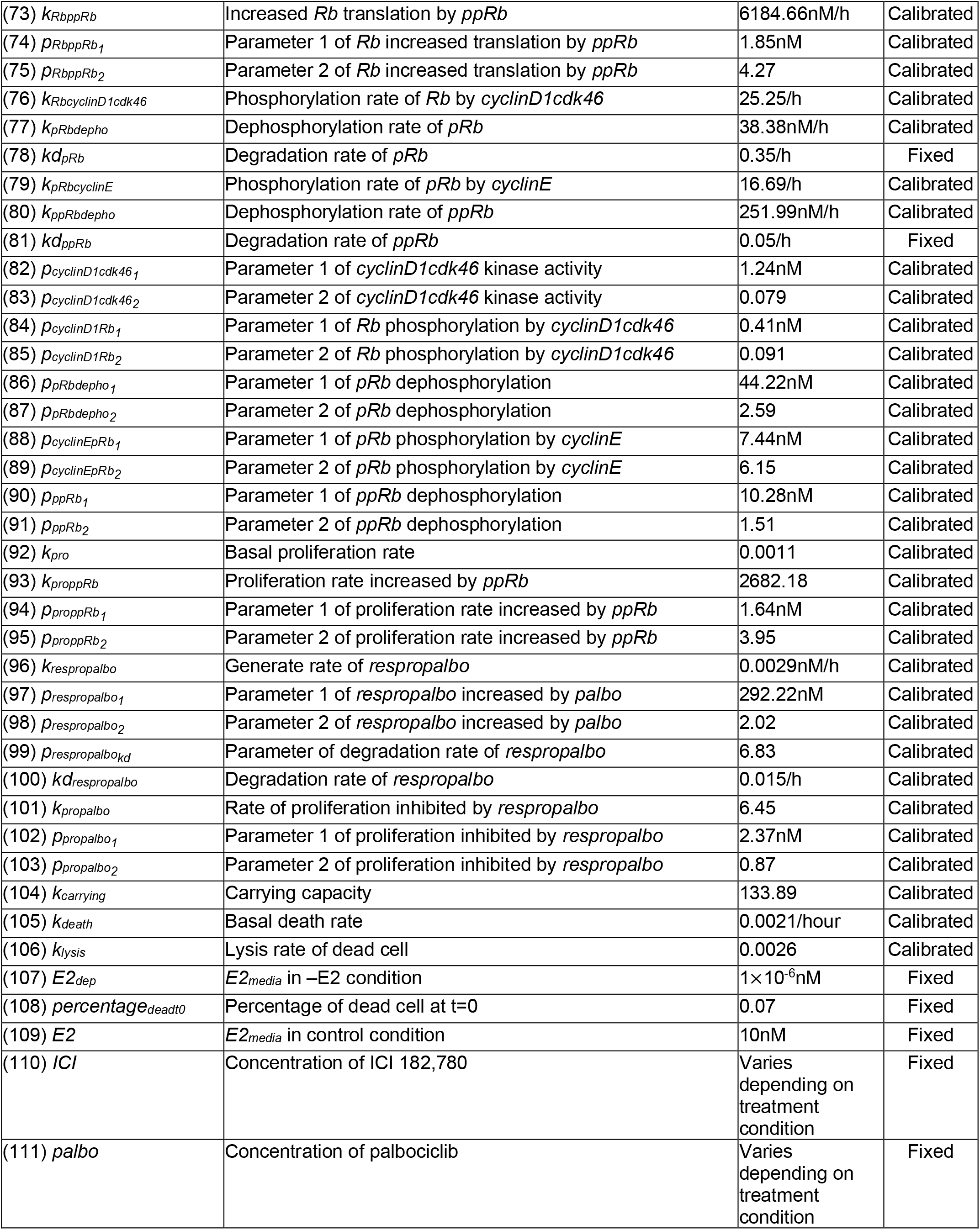
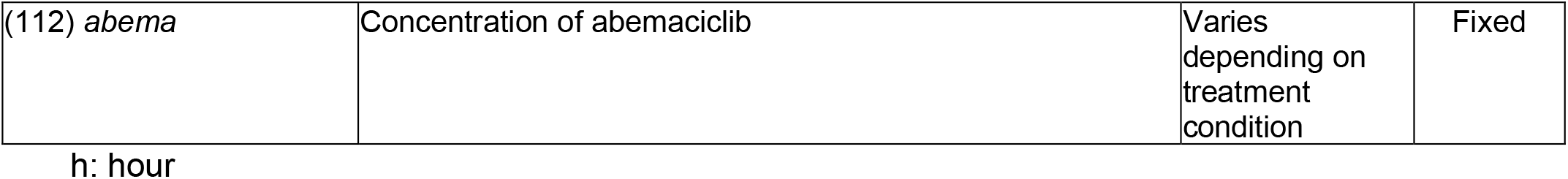
Model Parameter Descriptions, Values and Declaration of Fixed or Calibrated. Parameter names and model variable names are italicized.

### Mathematical Model

#### Biological Signaling Diagram

The structure of our ODE model is based on the signaling pathways of the G1-S transition since the drugs of interest, anti-estrogens and Cdk4/6 inhibitors, primarily affect progression through the G1 phase of the cell cycle (Musgrove, et al., 2009). The justification of the numbered interactions of the signaling pathway and drugs, shown in Figure 1A, is as follows:

1. –E2 decreases the level of estrogen (Lewis et al., 2005); 2. E2 binds to ER and forms the transcription factor E2:ER (Vrtačnik, et al., 2014); 3. ICI binds to ER and forms ICI:ER, which increases the degradation of ER and blocks its transcriptional activity (Xi et al., 2020); 4. E2:ER increases transcription of c-Myc (Prall et al., 1998); 5. E2:ER increases transcription of cyclinD1 (Prall et al., 1998); 6. E2:ER increases transcription of cyclinE (Musgrove, et al., 2009); 7. c-Myc inhibits transcription of p21 (Bretones et al., 2015); 8. CyclinD1 binds to Cdk4/6 and forms the cyclinD1:Cdk4/6 kinase (Sherr 1995); 9. CyclinE binds to Cdk2 and forms the cyclinE:Cdk2 kinase (Prall et al., 1998); 10. p21 binds to cyclinD1:Cdk4/6 and forms the cyclinD1:Cdk4/6:p21 complex, which inhibits its kinase activity (Sherr and Roberts, 1995); 11. p21 binds to cyclinE:Cdk2 and forms the cyclinE:Cdk2:p21 complex which inhibits its kinase activity (Musgrove et al,. 2011); 12. Palbociclib binds to Cdk4/6 and inactivates its activity (Wells et al., 2020); 13. Abemaciclib binds to Cdk4/6 and inactivates its activity (Wells et al., 2020); 14. Palbociclib binds to cyclinD1:Cdk4/6 and forms the cyclinD1:Cdk4/6:palbociclib complex, which inactivates its kinase activity (Well et al., 2020); 15. Abemaciclib binds to cyclinD1:Cdk4/6 and forms the cyclinD1:Cdk4/6:abemaciclib complex, which inactivates its kinase activity (Well et al., 2020); 16. p21 binds to cyclinD1:Cdk4/6:palbociclib and forms the cyclinD1:Cdk4/6:p21:palbociclib complex (Pack et al., 2021); 17. p21 binds to cyclinD1:Cdk4/6:abemaciclib and forms the cyclinD1:Cdk4/6:p21:abemaciclib complex (Pack et al., 2021); 18. Palbociclib binds to cyclinD1:Cdk4/6:p21 and forms the cyclinD1:Cdk4/6:p21:palbociclib complex (Pack et al., 2021); 19. Abemaciclib binds to cyclinD1:Cdk4/6:p21 and forms the cyclinD1:Cdk4/6:p21:abemaciclib complex (Pack et al., 2021); 20. CyclinD1:Cdk4/6 phosphorylates RB1 to RB1-p (hypophosphorylated RB1) (Prall et al., 1998); 21. CyclinE:Cdk2 phosphorylates RB1-p to RB1-pp (hyperphosphorylated RB1) (Prall et al., 1998); 22. RB1 binds to E2F and inhibits its transcriptional activity (Bretones et al., 2015); 23. RB1-p binds to E2F and inhibits it transcriptional activity (Bretones et al., 2015); 24. E2F up-regulates RB1 expression (Yao et al., 2011); 25. E2F up-regulates c-Myc expression (Álvaro-Blanco et al., 2009); 26. E2F up-regulates cyclinE expression (Morris et al., 2000); 27. E2F drives the G1-S cell cycle transition and proliferation (Stevens et al., 2003); 28. Cell death.

Treatment with Cdk4/6 inhibitors will affect the stability of Cdk4/6 complexes bound to Cdk Interacting Protein/Kinase Inhibitory Protein (CIP/KIP) protein inhibitors (p21). The Cdk4/6 inhibitors can dissociate p21 selectively from Cdk4 but not Cdk6 (Pack et al., 2020). Because we didn’t differentiate between Cdk4 and Cdk6, we didn’t exclude the possibility of forming the tetramers cyclinD1:Cdk4/6:p21:palbociclib and cyclinD1:Cdk4/6:p21:abemaciclib. And the degradation rate of cyclinD1:Cdk4/6:p21:palbociclib and cyclinD1:Cdk4/6:p21:abemaciclib are assumed to be the same as the degradation rate of cyclinD1:Cdk4/6:p21 trimer. The binding interactions between cyclinD1, Cdk4/6, p21, palbociclib and abemaciclib can form different dimers, trimers and tetramers, depending on the subtypes of the Cdks and CIPs/KIPs (Guiley et al., 2019; Pack et al., 2020). Because both Cdk4/6:palbociclib (Cdk4/6:abemaciclib) and cyclinD1:Cdk4/6:palbociclib (cyclinD1:Cdk4/6:abemaciclib) lose their kinase activity, we didn’t include the binding reaction between Cdk4/6:palbociclib (Cdk4/6:abemaciclib) and cyclinD1. We only use the binding reactions between palbociclib (abemaciclib) and cyclinD1:Cdk4/6 to form the cyclinD1:Cdk4/6:palbociclib (cyclinD1:Cdk4/6:abemaciclib) complex. And we didn’t include the inhibition potency of abemaciclib on Cdk2 and cyclinE:Cdk2 (Wells et al., 2020) since they are not needed to capture the abemaciclib treatment effects. For conciseness, lines with arrowheads representing the unbinding reactions corresponding to each binding reaction included in the model are not shown in Figures 1A and 1B.

#### Model Structure

Protein level changes in response to estrogen signaling and drug treatments are affected by thousands of interactions among proteins. Even though the interactions in Figure 1A are limited to the G1 phase of the cell cycle, the reactions shown are incomplete and many interactions at the G1-S phase transition are excluded. For example, in addition to RB1 and RB1-p, other pocket protein members p107 and p130 also bind to E2F and inhibit its transcriptional activity (Yao et al., 2011). And CyclinD1:Cdk4/6 can phosphorylate p107 and p130 and increase their degradation (Leng et al., 2002 and Tedesco 2002). Also, E2F up-regulates the expression of itself (Yao et al., 2011) and Cdc25A (Stevens et al., 2003), which is a protein phosphatase that removes the inhibitory phosphorylation on Cdk4/6 and Cdk2, positively regulating their kinase activities (Shen et al., 2012). It is impractical to include all possible reactions related to treatments in the biological mechanism. Because our goal is to build a model that can predict treatment responses over long time scales, we simplified the interactions shown in Figure 1A to those necessary to capture the effects of different treatments. The model structure we used is shown in Figure 1B and is modified from Figure 1A. First, we ignored interaction 26, E2F up-regulates cyclinE expression, in Figure 1A as this simplification doesn’t affect the model simulation results. Second, we didn’t include Cdk2 explicitly in the model but assumed that cyclinE not bound to p21 is bound to Cdk2 and active. This is because of the long-held presumption that Cdks are in excess of the cyclins in the cell and Cdk2 has been shown to be in excess of its cyclin partners (Arooz, et al., 2000). Last, we didn’t include E2F in the model but assume that the level of RB1-pp reflects the transcriptional activity of E2F. While E2F, as the last driver of G1-S transition, may be the best protein to govern the proliferation rate (Stevens et al., 2003), the situation is complicated. Considering there are six E2F family members, having divergent roles as transcriptional activators or inhibitors, and the complexity of all possible combinations of E2Fs with their partners (Cam et al., 2003), it is difficult to use one measured protein level to denote E2F transcriptional activity. In order to govern the proliferation rate by one protein level and measure its level to calibrate the model, we decide to use hyperphosphorylated RB1 (RB1-pp) to represent E2F transcriptional activity and govern the proliferation rate. The is because E2Fs as transcriptional activators are regulated principally through binding to RB1 (MacDonald et al., 2012) and are only released to transactivate the genes required for the G1-S transition when RB1 is fully inactivated after phosphorylation by cyclinD1:Cdk4/6 and cyclinE:Cdk2 (Bretones et al., 2015). These facts make it possible to drive the proliferation rate by one protein with a single specific phosphorylation site representing RB1-pp. It has been demonstrated that RB1 exists mainly in unphosphorylated, monophosphorylated and hyperphosphorylated form and measuring a specific phosphorylation site on RB1 can be used to infer the hyperphosphorylated state of RB1 (Chung et al., 2019). Over fifteen phosphorylation sites are found on RB1 and we found that RB1 phosphorylated on S612 reflects the decreased RB1-pp level changes after treatments based on our experiment results and the literature (MacDonald et al., 2012). Therefore, the binding reactions 22 and 23 in Figure 1A are ignored and the arrows of interactions 24, 25 and 27 start from RB1-pp in Figure 1B instead of E2F in Figure 1A. The other numbered interactions shown in Figure 1B are the same as in Figure 1A.

#### Long term palbociclib treatment effect on proliferation and cyclinD1

Figure S4 re-plots the 10-week palbociclib monotreatment data from Figure 4. MCF7 cells are treated with 750nM palbociclib for 10 weeks and the cells are re-plated at 5 weeks. The blue line is the cell number from 0 to 5 weeks normalized to the initial cell number at t=0. The red line is the re-plated cell number from 5 to 10 weeks normalized to the initial re-plated cell number at 5 weeks. The plot shows that the MCF7 proliferation rate significantly decreased from 5 weeks to 10 weeks compared to 0 to 5 weeks. In order to make the model match these proliferation changes, we introduced another variable in the model to control proliferation under palbociclib treatment *(respropalbo* in Table 1), which will increase under palbociclib treatment and decrease after removal of palbociclib. The proliferation rate is divided by a Hill function of this variable and will decrease as palbociclib treatment time increases. Without this variable, the model’s simulation of proliferation rate over the entire 10 week period will be nearly constant, which would be a poor fit to the experiment results. So, adding the variable is necessary to make the model match the proliferation difference shown in the experiment.

Figure 4E shows that the cyclinD1 level increased after mono or alternating palbociclib treatment, which is consistent with the literature (Cornell et al., 2019 and Pancholi et al., 2020). In order to allow the model to simulate the increase of cyclinD1, we introduce another variable in the model *(rescyclinD1palbo* in Table 1) which will increase under the palbociclib treatment and decrease after removal of palbociclib. The generation rate of cyclinD1 is added to a hill function of this variable and will increase with palbociclib treatment time. Adding the variable is necessary to make the model match the cyclinD1 increase shown in the experiment. Although the cyclinD1 level increased after mono or alternating palbociclib treatment, the proliferation rate of the MCF7 cells remained suppressed under treatment. In our model, the increase of cyclinD1 can not counterbalance the palbociclib treatment effect of decreasing the cyclinD1:Cdk4/6 level. Although the increase of cyclinD1 causes the cyclinD1:Cdk4/6 level to rebound after the initial sharp decrease following the start palbociclib treatment, its level is still lower than the level before palbociclib treatment and the phosphorylation of RB1 by cyclinD1:Cdk4/6 is decreased. We model the phosphorylation rate of RB1 by cyclinD1:Cdk4/6 as a hill function of cyclinD1:Cdk4/6 multiplying a hill function of RB1, instead of cyclinD1:Cdk4/6 multiplying a hill function of RB1.

This modification allows us to better control the phosphorylation rate of RB1 by cyclinD1:Cdk4/6 in the model to match the decreased proliferation and increased cyclinD1 level under the palbociclib treatments.

#### Model Equations

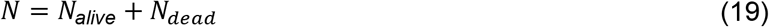

(19) Total number of cells equals number of alive cells plus number of dead cells

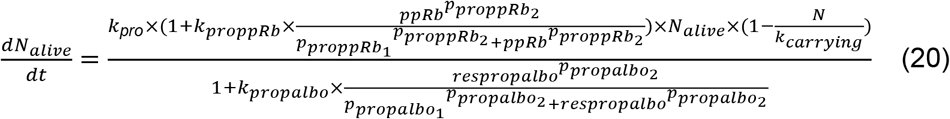

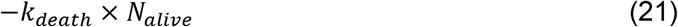

(20) Basal proliferation, increased proliferation by *ppRb* and inhibited proliferation by *respropalbo*

(21) Basal death

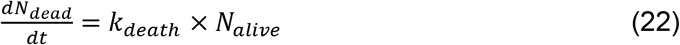

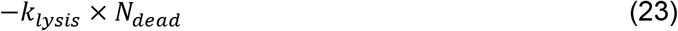

(22) Basal death

(23) Lysis of dead cells

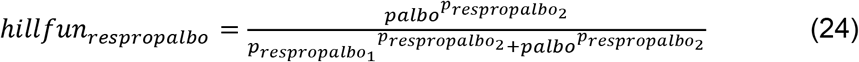

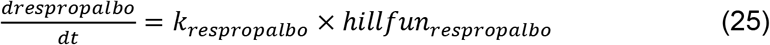

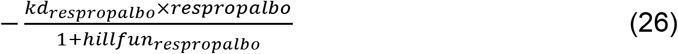

(24) Hill function for *respropalbo*

(25) Generation of *respropalbo* by *palbo*

(26) Degradation of *respropalbo* (fast if no palbociclib, but slow if there is palbociclib to allow slow buildup of *respropalbo*)

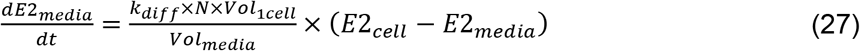

(27) E2 concentration changes in media

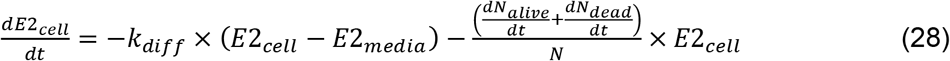

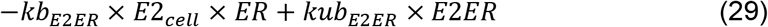

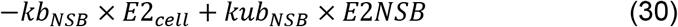

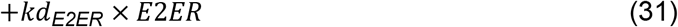

(28) E2 concentration changes in cell

(29) Binding and unbinding between *ER* and *E2_cell_*

(30) Binding and unbinding between non-specific binding and *E2_cell_* in the cell

(31) Degradation of *E2ER*

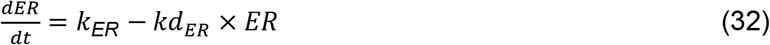

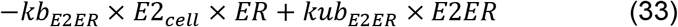

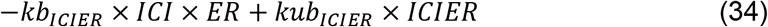

(32) Translation and degradation of *ER*

(33) Binding and unbinding between *ER* and *E2_cell_*

(34) Binding and unbinding between *ER* and *ICI*

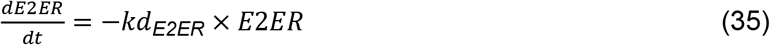

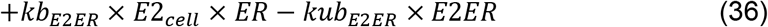

(35) Degradation of *E2ER*

(36) Binding and unbinding between *ER* and *E2_cell_*

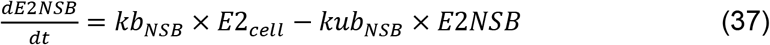

(37) Binding and unbinding between non-specific binding and *E2_cell_*

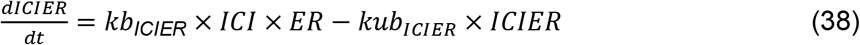

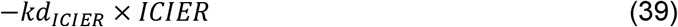

(38) Binding and unbinding between *ICI* and *ER*

(39) Degradation of *ICIER*

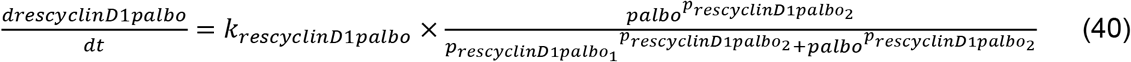

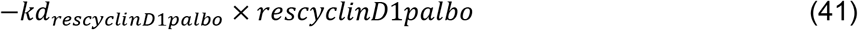

(40) Generation of *rescyclinD*1*palbo* by *palbo*

(41) Degradation of *rescyclinD*1*palbo*

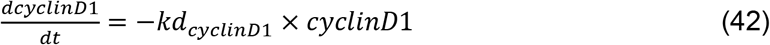

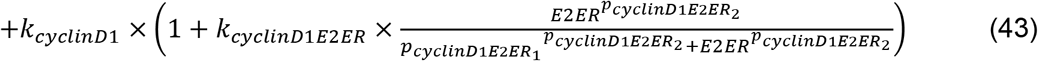

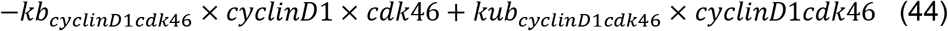

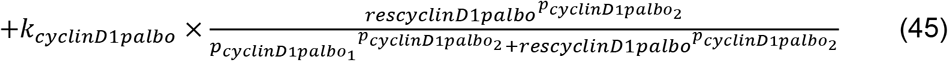

(42) Degradation of *cyclinD1*

(43) Basal translation of *cyclinD1* and the increased translation by *E2ER*

(44) Binding and unbinding between *cyclinD1* and *cdk46*

(45) Increased translation of *cyclinD1* by *rescyclinD*1*palbo*

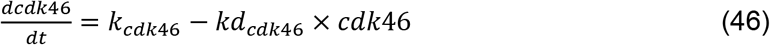

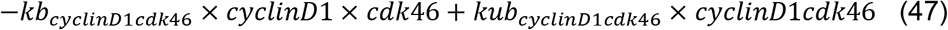

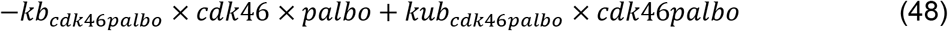

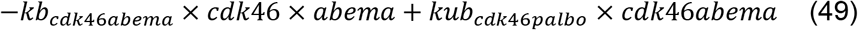

(46) Translation and degradation of *cdk46*

(47) Binding and unbinding between *cyclinD1* and *cdk46*

(48) Binding and unbinding between *palbo* and *cdk46*

(49) Binding and unbinding between *abema* and *cdk46*

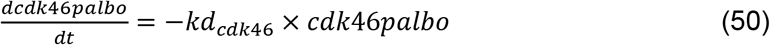

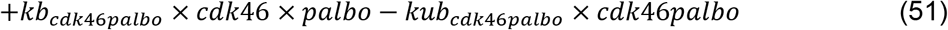

(50) Degradation of *cdk46palbo*

(51) Binding and unbinding between *palbo* and *cdk46*

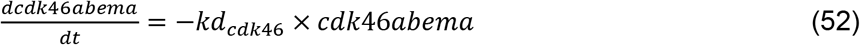

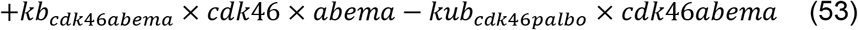

(52) Degradation of *cdk46abema*

(53) Binding and unbinding between *abema* and *cdk46*

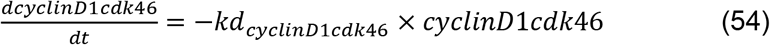

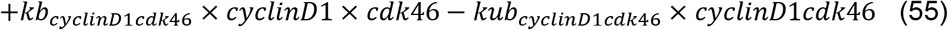

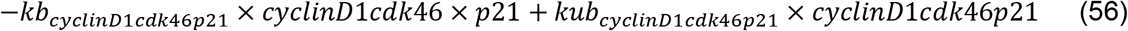

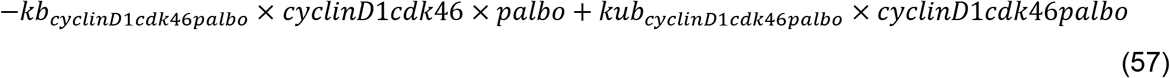

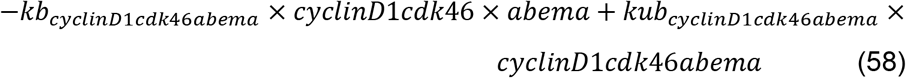

(54) Degradation of *cyclinD*1*cdk46*

(55) Binding and unbinding between *cyclinD1* and *cdk46*

(56) Binding and unbinding between *p*21 and *cyclinD*1*cdk46*

(57) Binding between *palbo* and *cyclinD*1*cdk46*

(58) Binding between *abema* and *cyclinD*1*cdk46*

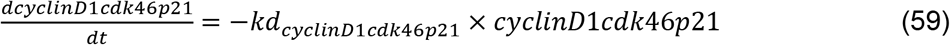

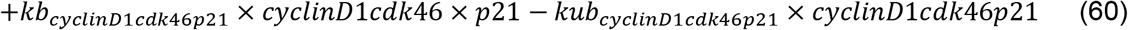

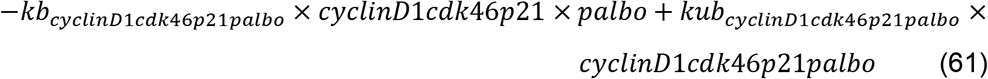

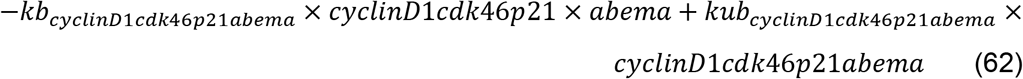

(59) Degradation of *cyclinD*1*cdk46p21*

(60) Binding and unbinding between p21 and *cyclinD*1*cdk46*

(61) Binding and unbinding between *palbo* and *cyclinD*1*cdk46p21*

(62) Binding and unbinding between *abema* and *cyclinD*1*cdk46p21*

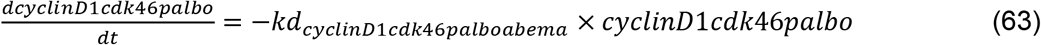

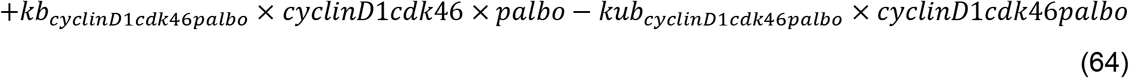

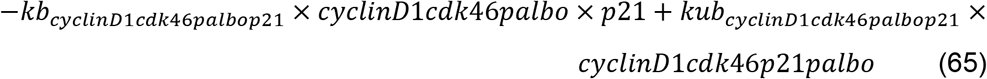

(63) Degradation of *cyclinD1cdk46palbo*

(64) Binding and unbinding between *palbo* and *cyclinD*1*cdk46*

(65) Binding and unbinding between *p*21 and *cyclinD*1*cdk*46*palbo*

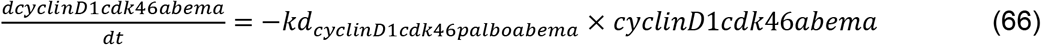

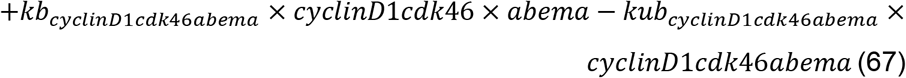

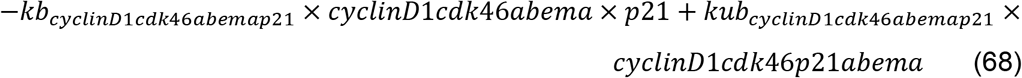

(66) Degradation of *cyclinD1cdk46abema*

(67) Binding and unbinding between *abema* and *cyclinD*1*cdk46*

(68) Binding and unbinding between *p*21 and *cyclinD1cdk46abema*

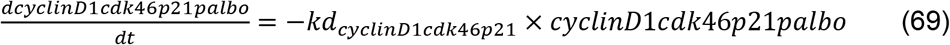

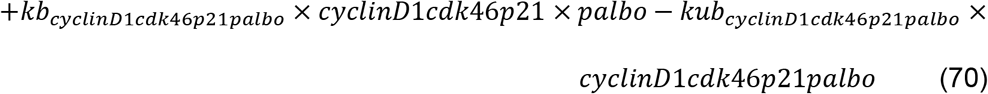

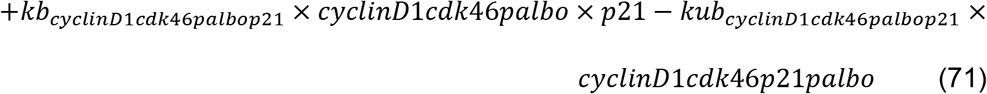

(69) Degradation of *cyclinD1cdk46p21palbo*

(70) Binding and unbinding between *palbo* and *cyclinD*1*cdk*46*p*21

(71) Binding and unbinding between *p*21 and *cyclinD*1*cdk*46*palbo*

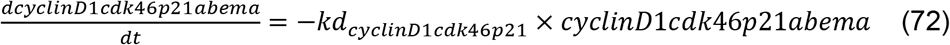

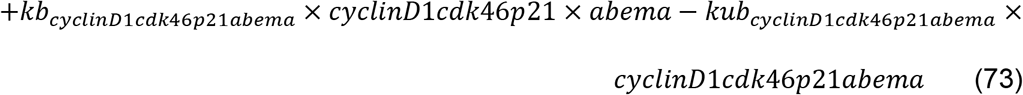

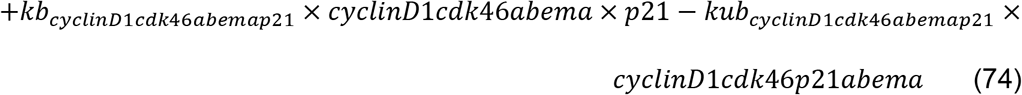

(72) Degradation of *cyclinD*1*cdk*46*p*21*abema*

(73) Binding and unbinding between *abema* and *cyclinD*1*cdk*46*p*21

(74) Binding and unbinding between *p*21 and *cyclinD*1*cdk*46*abema*

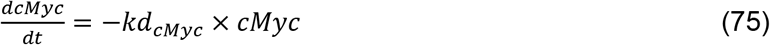

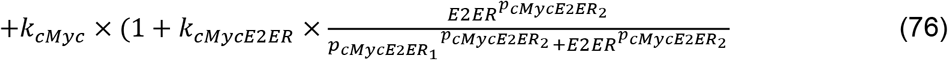

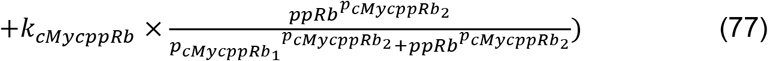

(75) Degradation of *cMyc*

(76) Basal translation of *cMyc* and the increased translation by *E2ER*

(77) Increased translation of *cMyc* by *ppRb*

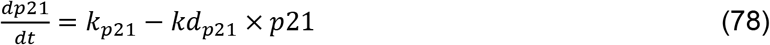

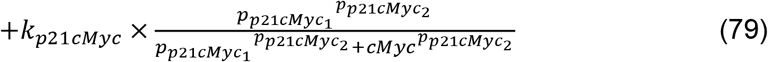

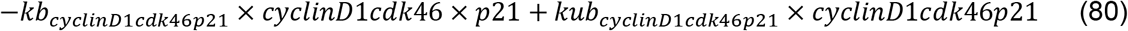

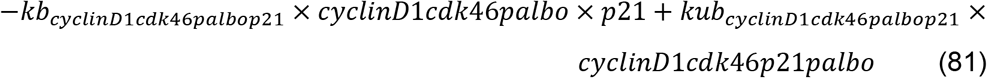

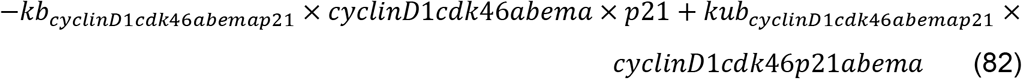

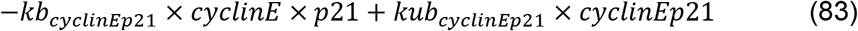

(78) Translation and degradation of *p*21

(79) Inhibition of translation by *cMyc*

(80) Binding and unbinding between *p*21 and *cyclinD*1*cdk46*

(81) Binding and unbinding between *p*21 and *cyclinD*1*cdk*46*palbo*

(82) Binding and unbinding between *p*21 and *cyclinD*1*cdk*46*abema*

(83) Binding and unbinding between *p*21 and *cyclinE*

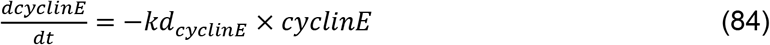

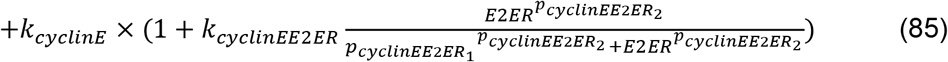

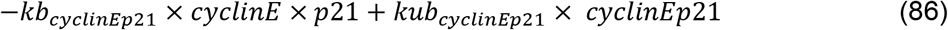

(84) Degradation of *cyclinE*

(85) Basal translation of *cyclinE* and the increased translation by *E2ER*

(86) Binding and unbinding between *p*21 and *cyclinE*

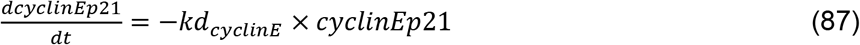

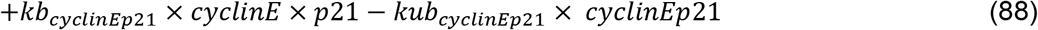

(87) Degradation of *cyclinEp*21

(88) Binding and unbinding between *p*21 and *cyclinE*

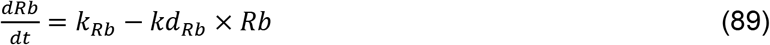

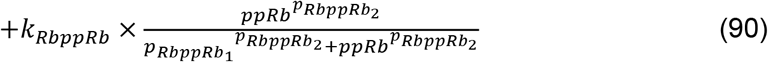

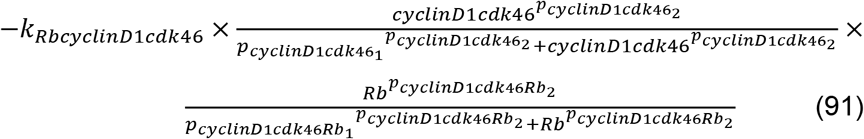

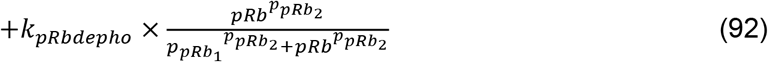

(89) Degradation and basal translation of *Rb*

(90) Increased translation of *Rb* by *ppRb*

(91) Phosphorylation of *Rb* by *cyclinD*1*cdk*46

(92) Dephosphorylation of *pRb*

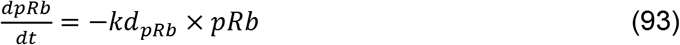

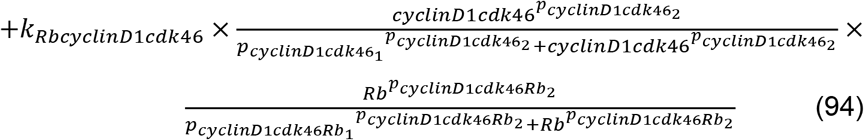

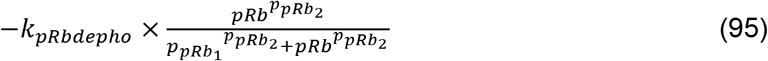

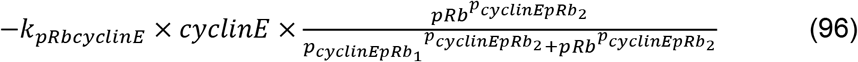

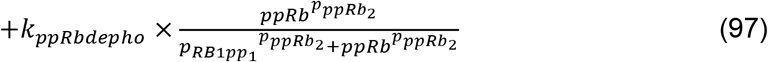

(93) Degradation of *pRb*

(94) Phosphorylation of *Rb* by *cyclinD*1*cdk*46

(95) Dephosphorylation of *pRb*

(96) Phosphorylation of *pRb* by *cyclinE*

(97) Dephosphorylation of *ppRb*

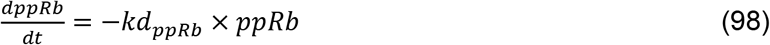

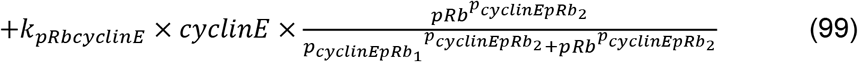

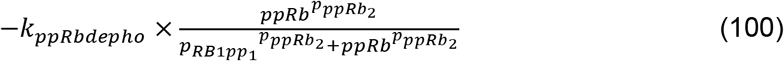

(98) Degradation of *ppRb*

(99) Phosphorylation of *pRb* by *cyclinE*

(100) Dephosphorylation of *ppRb*

### Model Parameter Calibration and Model Summary

Parameters of degradation rates (*k_d_*) for proteins were assigned according to half-lives found in literature (denoted as Fixed in Table 2), where *k_d_* = −log(1/2)/t_half-life_. Considering that the limited data we collected does not warrant the increase of parameter number, which would be unidentifiable, we facilitated the optimization and decreased the number of parameters to be calibrated by fixing the unbinding parameters to 1 (denoted as Fixed in Table 2). The other parameters were calibrated using the *patternsearch* function in MATLAB (R2021b), to minimize the discrepancy between the model simulation and the experimental results. The least-squares cost function was calculated as:

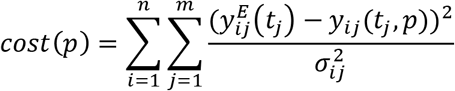

where *i* indexes the model variables of proteins or alive cell number under a specific treatment, *j* indexes the experimental measured times. 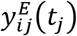 is the experimental measurement of the specie *i* at time *j*. *y_ij_*(*t_j_, p*) is the simulation result of the variable *i* at time *j* using parameter vector *p. σ_ij_* is the standard deviation of the experimentally measured specie *i* at time *j*. The data, time points and the model variables used in fitting the experimental data are listed in Table S1. The parameters were tuned manually at the beginning to get the gross treatment response roughly consistent with the experimental results. Then the default *patternsearch* in MATLAB was used for calibration of the parameters.

The culture media, including any drugs in the different treatments, was changed at t = 0 and every 3 or 4 days during the treatment period, so the longest period without resupply of any drug is 4 days. There is no data we are aware of for the half-life of the three drugs we used (ICI 182,780, palbociclib and abemaciclib) in in-vitro conditions. Most data are for the plasma or terminal half-life in-vivo determined by processing in the liver or excretion through the kidneys, which are not applicable to our model. There is data for the in-vitro stability of these drugs in human plasma, which is akin to our case. ICI 182,780 shows no degradation in human plasma at room temperature over 7 hours (Alegete et al., 2017). Palbociclib and abemaciclib show less than 5% degradation in human plasma at room temperature over 3 days (Martinez-Chavez et al., 2019). Based on this data, we believe the drug half-life is sufficiently long so as to be ignored in our in-vitro experiments. The drug concentrations are assumed to be constant throughout the treatment as a reasonable approximation.

The mathematical model contains 27 ordinary differential equations (ODEs) and 112 parameters (80 calibrated and 32 fixed), which is implemented in MATLAB. The generation, degradation, phosphorylation, dephosphorylation, binding and unbinding reactions are modeled by mass action laws and hill functions. Drug treatment effects are modeled by competitive binding to their targets. The ODEs are solved numerically by the *ode23tb* function in MATLAB.

### Parameter cohort

To address the fact that the parameters in model may not be practically identifiable by our limited measured data, another 199 parameter sets that fit the data reasonably well (cost < 500) were identified to form a parameter cohort. All 200 parameter sets in the cohort were used for simulation and prediction. The resulting spread in the predictions show the degree to which the data used to calibrate the model parameters constrains the prediction results. The parameter cohort was generated by the default genetic algorithm function, *ga*, in the MATLAB optimization toolbox. We saved parameter sets found during the running of the *ga* function whose cost function value was smaller than 500.

The coefficients of variation of the parameters and the cost function values for the parameter sets in the cohort are plotted in Figures S9A and S9B. The coefficient of variation plot reflects the spread of the parameter values in the cohort and the large values represent parameters to which the model has low sensitivity. A local sensitivity analysis for each parameter in the cohort with respect to proliferation is shown in Figure S9D. The most significant sensitivity for cell proliferation involves parameters #6 and #90, which are related to the basal translation of the estrogen receptor and the dephosphorylation of RB1-pp, respectively. These results are not surprising as the estrogen receptor level impacts the response to –E2 and ICI endocrine treatments and dephosphorylation of RB1-pp directly regulates the RB1-pp level. Figure S9C plots a histogram of the fitting costs that were generated by all the perturbed parameter sets used in the sensitivity analysis used to create Figure. S9D. The figure shows that across all the perturbations of parameter sets in the cohort there was not a major increase in the fitting cost.

### Local Sensitivity Analysis

We used local sensitivity analysis to check how sensitive the model output was to the parameter values. The sensitivity value for a model output and a specific parameter is the change in the model output relative to the change in parameter value. It can be expressed as (Zi et al., 2011 and Nagaraja et al., 2014):

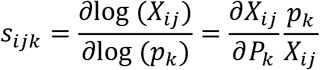

where *s_ijk_* is the local sensitivity value, which is the derivative of model output *X_ij_* with respect to parameter *p_k_* multiplied by the ratio *p_k_/X_ij_*. It gives the percent change in the model output produced by a 1%) change in a parameter. In the equation, *i* indexes the alive cell number under a specific treatment, *j* indexes the timepoints and *k* indexes the parameters. *s_ijk_* can be approximated by the second order central finite difference. Therefore, each parameter is individually varied by 1% of its value:

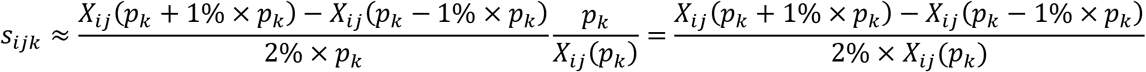

The sensitivity analysis was performed on all 64 data-calibrated parameters in a parameter set, except the hill function powers, and all parameter sets in the cohort. The fixed parameters were excluded. Because we want to check whether the model output is very sensitive to certain parameters, the maximum values of *s_ijk_* across all *i* and *j*, which is 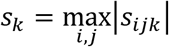, is used to represent the sensitivity value.

### Growth Rate Inhibition (GR) Metric

The GR metric is different from traditional drug response metrics, which are highly sensitive to the number of cell divisions during the experiment. It compares the growth rates in the presence and absence of drug and is largely independent of cell division rate and assay duration (Hafner et al., 2016). The GR metric is calculated according to the formula:

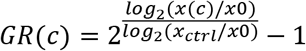

The cell count under the drug treatment is normalized to the vehicle control cell count. *x*(*c*) is the cell count in the presence of drug at concentration *c*. *x_ctrl_* is the cell count for the vehicle control. *x*_0_ is the cell count at t = 0 prior to drug treatment. The maximum value of GR is 1 (unless the drug increases proliferation) and the lowest value of GR is −1. GR = 0 means the drug treatment has a cytostatic response and a negative value means the drug treatment has a cytotoxic effect (Hafner et al., 2016).

### Microarray Data Analysis

The microarray data analysis was performed by the limma R package, which provides data normalization and differential gene expression analysis for gene expression experiments (Ritchie et al., 2015). The agilent microarray data files were read by the *read.maimages* function. The backgrounds were corrected using the *backgroundCorrect* function and the data were normalized using the *normalizedBetweenArrays* function. Differential expression analysis was performed by *lmfit* and *eBayes* functions. Heatmaps of the significantly differentially expressed genes (adjusted p-value <= 0.05) were plotted using the *heatmap.2* function in the gplots R package. Hierarchical clustering of columns in the heatmap is based on the default setting in the *heatmap.2* function, which used the *dist* and *hclust* functions in the stats R package. Principal component analysis was performed using the *prcomp* function in the stats R package. The Gene Set Enrichment Analysis (GSEA) was performed using the clusterProfiler R package (Yu et al., 2012 and Wu et al., 2021) and the C3 regulatory target gene sets in the Molecular Signatures Database (MSigDB) provided by the msigdbr R package was used (Subramanian et al., 2005). Data preparation and visualizations were performed in R using the tidyverse (v1.3.1), gplots (v3.1.1), ggplot2(3.3.5), and plotly(4.10.0) packages

### Statistical Analysis

Statistical testing was carried out in MATLAB. Group comparisons were performed by the two-sided paired t test (*ttest* function). For comparison of multiple groups, one-way ANOVA or two-way ANOVA (*anovan* function) and Tukey’s HSD test for multiple comparisons (*multcompare* function) was used. Lower case n refers to the number of biological replicates noted in the figure legends. Statistical significance was considered with p values smaller than 0.05 and ns represents non-significant. The precise p values are noted in the figure legends with asterisks: p <0.05 (*), p ≤ 0.01 (**), p ≤ 0.001 (***), p ≤ 0.0001 (****).

