## Supplemental file for "Modeling Breast Cancer Proliferation, Drug Synergies, and Alternating Therapies"

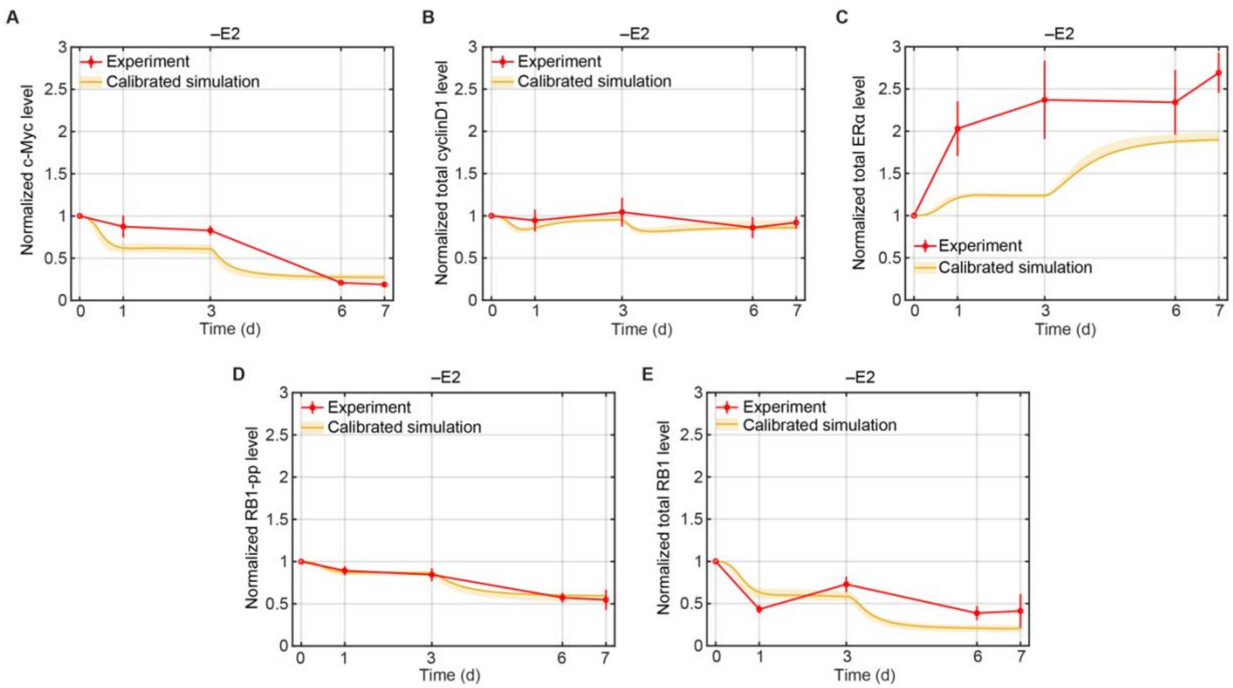


**Figure S1. Model Calibration Simulations of Normalized Protein Levels for –E2 Treatment Compared to Experimental Data.**

(A) Model simulation of normalized c-Myc compared to experimental data (mean ± s.e., n=3). The experimental data are shown in red and the simulation results are shown in yellow (solid line represents the lowest cost value simulation and the shaded regions contains the central 98% of the cohort simulations).

(B) Model simulation of normalized total cyclinD1 compared to experimental data (mean ± s.e., n=3). Color, lines and the shaded regions have the same meaning as (A).

(C) Model simulation of normalized total ERα compared to experimental data (mean ± s.e., n=3).

(D) Model simulation of normalized RB1-pp compared to experimental data (mean ± s.e., n=3).

(E) Model simulation of normalized total RB1 compared to experimental data (mean ± s.e., n=3). The experimental data are re-used from previous work (He et al., 2020).

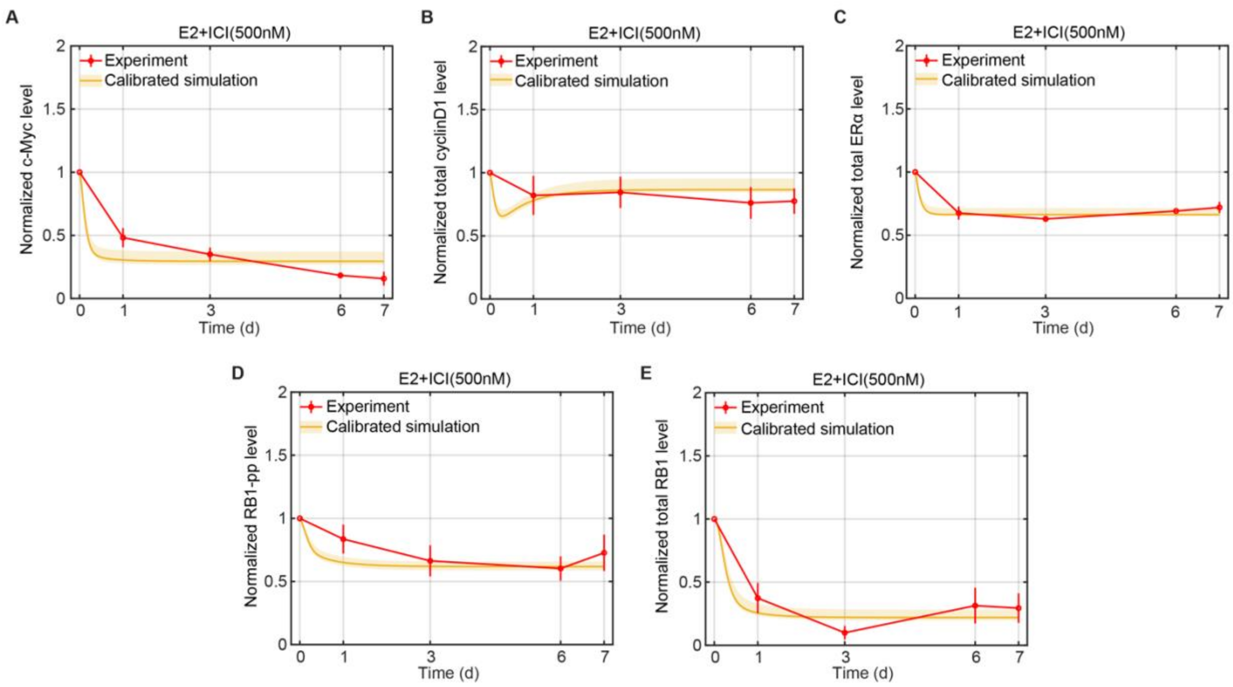


**Figure S1. Model Calibration Simulations of Normalized Protein Levels for E2+ICI(500nM) Treatment Compared to Experimental Data.**

(A) Model simulation of normalized c-Myc compared to experimental data (mean ± s.e., n=3). The experimental data are shown in red and the simulation results are shown in yellow (solid line represents the lowest cost value simulation and the shaded regions contains the central 98% of the cohort simulations).

(B) Model simulation of normalized total cyclinD1 compared to experimental data (mean ± s.e., n=3). Color, lines and the shaded regions have the same meaning as (A).

(C) Model simulation of normalized total Erα compared to experimental data (mean ± s.e., n=3).

(D) Model simulation of normalized RB1-pp compared to experimental data (mean ± s.e., n=3).

(E) Model simulation of normalized total RB1 compared to experimental data (mean ± s.e., n=3). The experimental data are re-used from previous work (He et al., 2020).

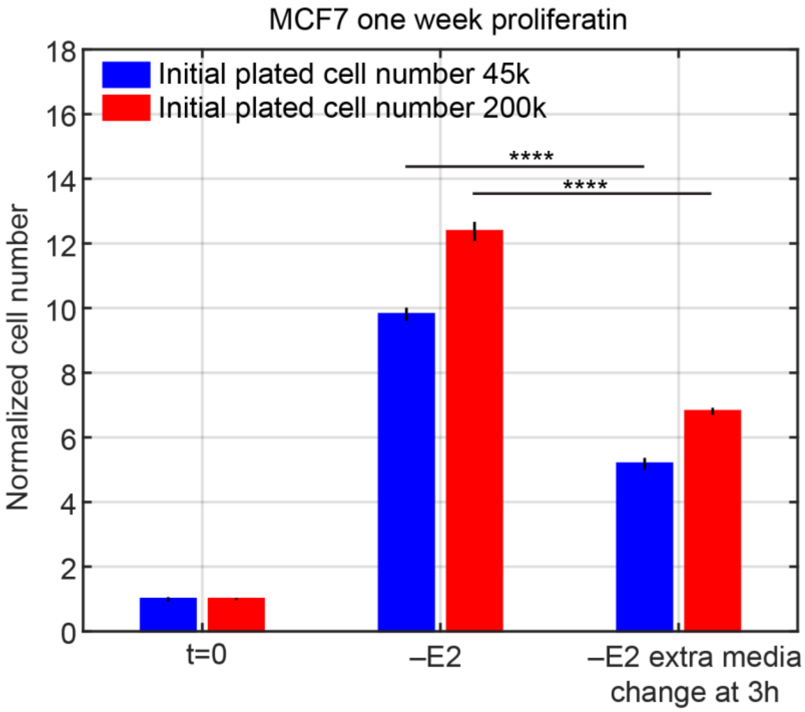


**Figure S3. Extra Media Change during –E2 Treatment Decreases MCF7 Cell Proliferation.** The bar plot shows normalized cell number relative to t=0 for the cases of $45\times10^3$(45k, blue bars) and $200\times10^3$(200k, red bars) initially plated cells (mean ± s.e., n=2 biological replicates, and 3 technical replicates for each biological replicate). The media is changed at t=0 and at day 3 to E2-deprived media in the –E2 case (middle). An extra media change to E2-deprived media at 3 hours was added in the –E2 extra media change case (right). In both cases of initially-plated cells, the extra media change, which further decreases the residual E2 level, significantly decreases the overall MCF7 proliferation at 1 week. Data were compared by two-way ANOVA, $p \leq 0.0001$ (****).

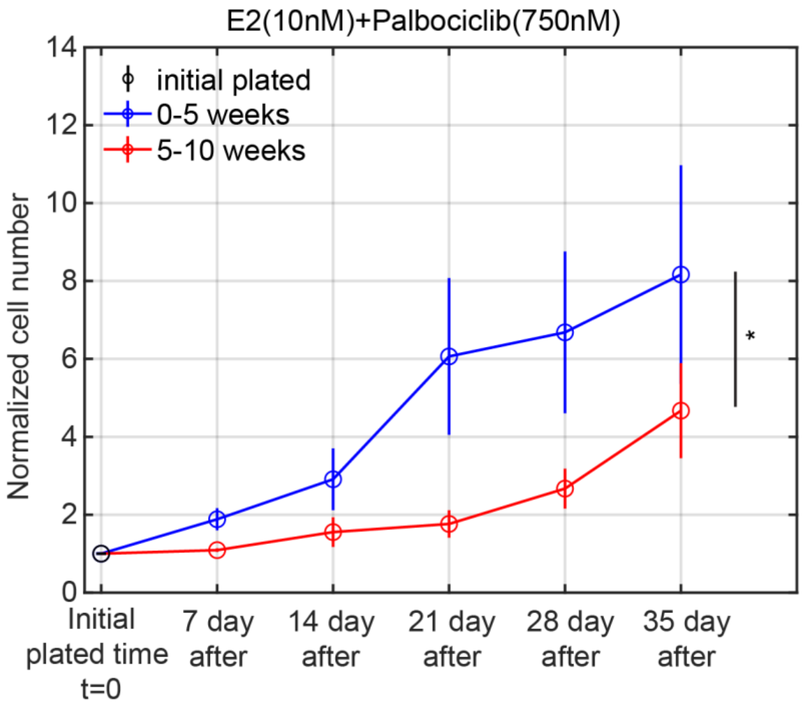


**Figure S4. MCF7 Cell Proliferation Rate Decreases from 5 Weeks to 10 weeks Compared to 0 to 5 Weeks in Palbociclib Treatment.** Plot shows normalized cell number relative to initial plated number under E2+palbo(750nM) treatment (mean ± s.e., n=3). This is a re-plot of the 10 weeks E2+palbo(750nM) treatment data in Figure 5. MCF7 cells were under E2+palbo (750nM) treatment for total 10 weeks. At 5 weeks, the cells were re-plated. The blue line is the normalized cell number from 0 to 5 weeks relative to the initial plated number at t=0. The red line is the normalized cell number from 5 to 10 weeks relative to the initial plated number at 5 weeks. The normalized proliferation from 5 to 10 weeks is significantly smaller than that from 0 to 5 weeks. Data were compared by paired t test, $p \leq 0.05$ (*).

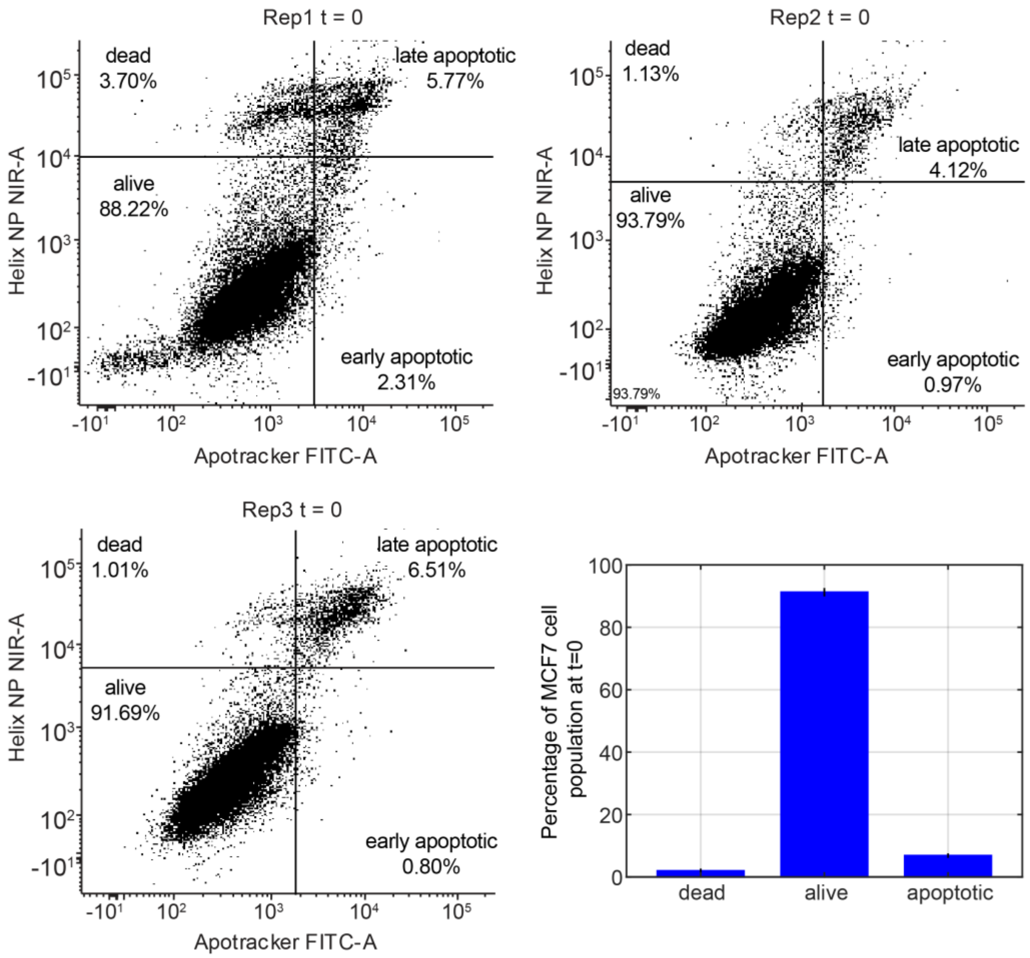


**Figure S5. Apoptosis Assay of MCF7 Cells at t=0.** Plots show the three replicate measurements for apoptotic percentage of MCF7 cells at t=0 by flow cytometry. The bar plot shows the average percentage of the three replications (mean ± s.e., n=3). The apoptotic percentage equals the early apoptotic percentage plus the late apoptotic percentage. The alive and apoptotic percentages were used to assign the *Nalive and Ndead* values in Table 1.

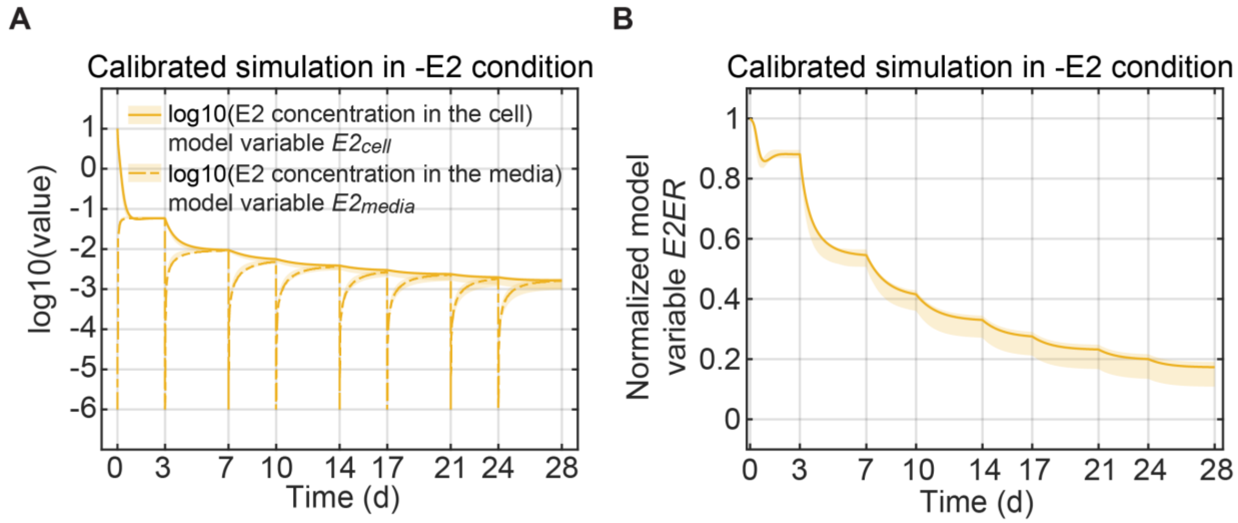


**Figure S6. Simulations of Model E2 Changes during –E2 Treatment.**

(A) Simulation of E2 concentration in cells (model variable $E2_{cell}$, number 2 in Table 1) and medium (model variable $E2_{media}$, number 1 in Table 1) during –E2 treatment. The solid and dashed lines represent the lowest cost value simulation and the shaded regions contain the central 98% of the cohort simulations.

(B) Simulation of model variable *E2ER* (number 4 in Table 1) during –E2 treatment. The lines and shaded regions have the same meaning as (A).

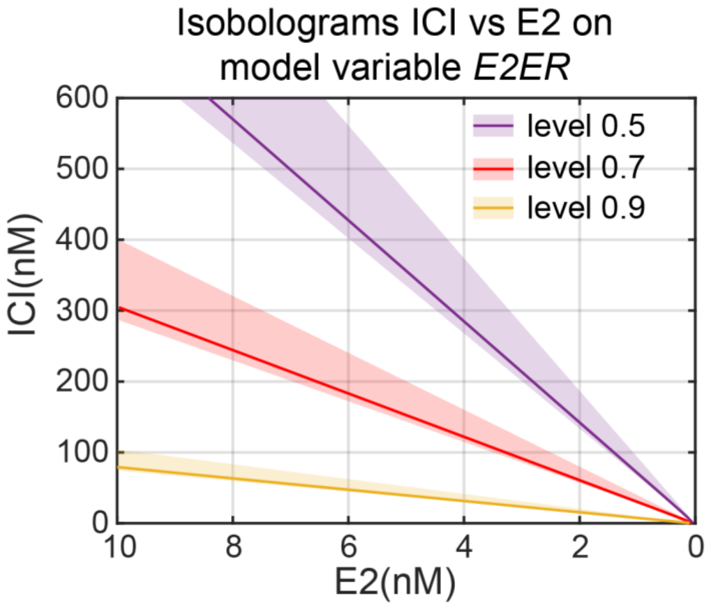


**Figure S7. Model simulation of isobologram between ICI and E2 on Normalized Model Variable *E2ER* at 17days.**

Different color of the isobole represents the different level of normalized model variable *E2ER*. The solid line representing the lowest cost value simulation and the shaded regions containing the central 98% of the cohort simulations.

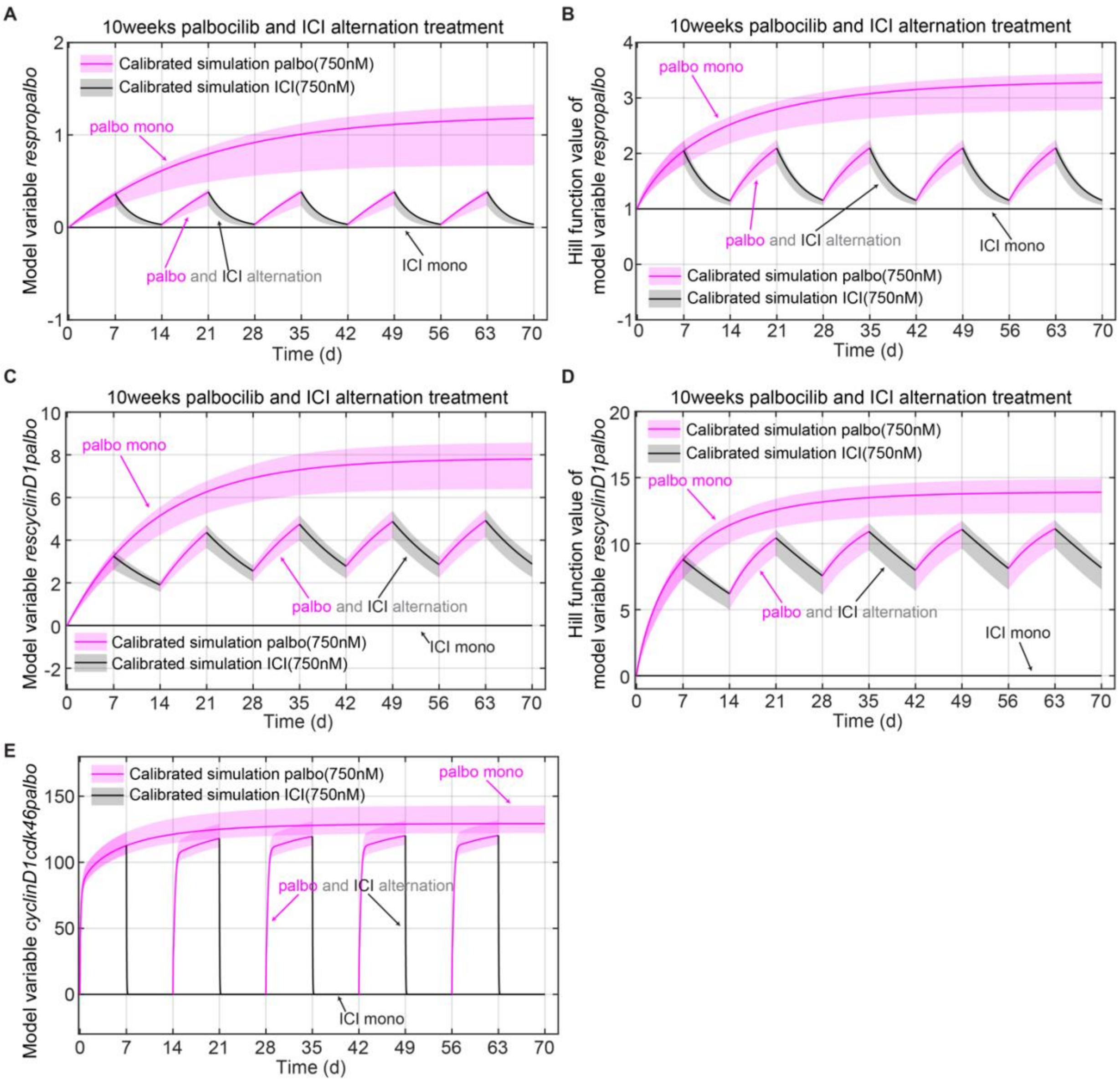


**Figure S8. Simulations of Model Variable Changes during Long Time Mono and Alternating Treatments.**

(A) Simulation of model variable *respropalbo* (number 25 in Table 1) changes during the mono and alternating treatments, which are E2+palbo(750nM), E2+ICI(750nM) and E2+palbo(750nM) alternating with E2+ICI(750nM) treatments, as in Figure 4A-4D. In both the mono and alternating treatments, the E2+palbo(750nM) condition is shown in purple and the E2+ICI(750nM) condition in black. In the alternating treatment, each treatment period is 7days and starts with

E2+palbo(750nM). The solid line represents the lowest cost value simulation and the shaded regions contain the central 98% of the cohort simulations.

(B) Simulation of changes in the hill function value in the model variable *respropalbo* (denominator of equation 20) during the mono and alternating treatments shown in (A). The lines and shaded regions have the same meaning as (A).

(C) Simulation of model variable *rescyclinD1palbo* (number 7 in Table 1) changes during the mono and alternating treatments shown in (A).

(D) Simulation of changes in the hill function value in the model variable *rescyclinD1palbo* (equation 45) during the mono and alternating treatments shown in (A).

(E) Simulation of model variable *cyclinD1cdk46palbo* (number 14 in Table 1) changes during the mono and alternating treatments shown in (A).

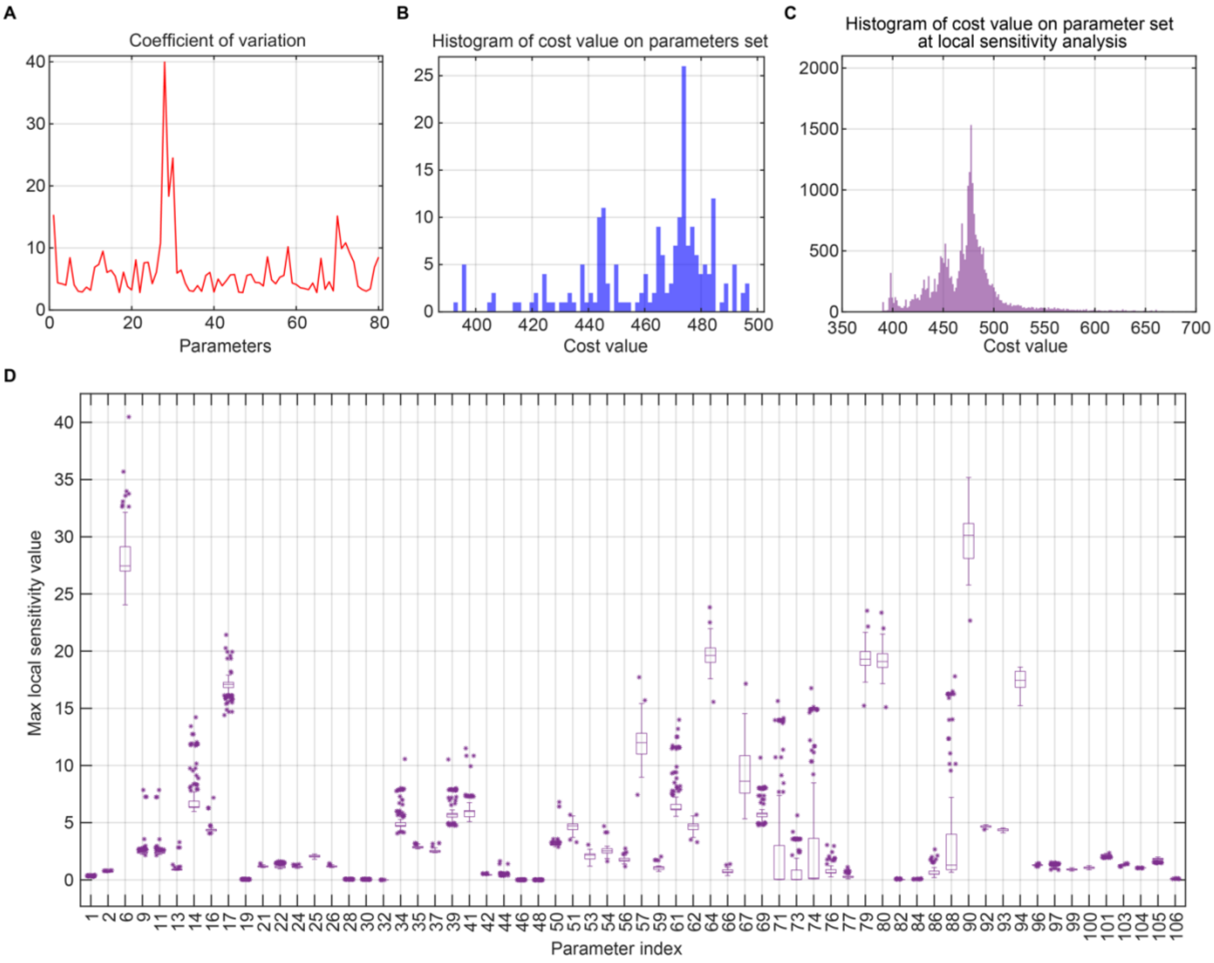


**Figure S9. Profile of the Parameter Sets in the Parameter Cohort.**

(A) Coefficients of variation of the parameters in the parameter cohort.

(B) Histogram of the cost values of each the parameter set in the parameter cohort.

(C) Boxplot of local sensitivity analysis of the model parameters. The parameter index is the index in Table 2. The analysis was performed on all 64 data-calibrated parameters in a parameter set, except hill function powers, and all parameter sets in the cohort. The maximum local sensitivity values across all proliferation data points for each parameter in the cohort are plotted in one box. The center line in each box is the median, and the bottom and top lines of each box are the 25$^{th}$ and 75$^{th}$ percentiles, respectively. The whiskers are maximum and minimum values without considering outliers. Data points are considered outliers if they lie more than 1.5×IQR (interquartile range) below the 25$^{th}$ percentile or above the 75$^{th}$ percentile.

(D) Histogram of the cost values for each perturbed parameter set used in the local sensitivity analysis.

Table S1. Times when various species were measured and used for parameter calibration or result prediction. E2 (control), E2+ICI (treated with ICI at a specific concentration in E2 medium), –E2 (E2 deprivation), –E2+ICI (treated with ICI at a specific concentration in –E2 medium), E2+palbo (treated with palbociclib at a specific concentration in E2 medium), E2+palbo+ICI (treated with palbociclib and ICI at a specific concentration in E2 medium),–E2+palbo (treated with palbociclib at a specific concentration in –E2 medium) and E2+abema (treated with abemaciclib at a specific concentration in E2 medium), E2+abema+ICI (treated with abemaciclib and ICI at a specific concentration in E2 medium). n: biological replications, d: days.

| Fi gu | Conditi on | Specie s name | Measure time point | Calibr ation/P | Model variables used for simulation |
|---|---|---|---|---|---|
| 1 C | E2 | normali zed cell [illegible] | 1d, 3d, 6d, 7d, 11d. Each time point n=3 | Calibr ation | *Nalive* |
| 1 D | –E2 | normali zed cell [illegible] | 1d, 3d, 6d, 7d, 14d, 21d, each time point n=3 | Calibr ation | *Nalive* |
| 1 E | E2+ICI (100nM [illegible] | normali zed cell [illegible] | 1d, 3d, 6d, 7d, 14d, 21d, each time point n=3 | Calibr ation | *Nalive* |
| 1F | E2+ICI (500nM [illegible] | normali zed cell [illegible] | 7d, 14d, 21d, each time point n=3 | Calibr ation | *Nalive* |
| 1 G | E2+pal bo | normali zed cell [illegible] | 7d, 14d, 21d, 28d each time point n=3 | Calibr ation | *Nalive* |
| 1 H | E2+pal bo | normali zed cell [illegible] | 7d, 14d, 21d, 28d each time point n=3 | Calibr ation | *Nalive* |
| 1I | E2+pal bo | normali zed cell [illegible] | 7d, 14d, 21d, each time point n=3 | Calibr ation | *Nalive* |
| 1J | – E2+ICI | normali zed cell [illegible] | 7d, 14d, 21d, each time point n=3 | Calibr ation | *Nalive* |
| 1 K | E2+pal bo | normali zed cell [illegible] | 7d, 14d, 21d, each time point n=3 | Calibr ation | *Nalive* |
| 2 A | E2+ab ema | normali zed cell [illegible] | 1d, 3d, 6d, 7d, 14d, 21d, each time point n=3 | Calibr ation | *Nalive* |
| 2 B | E2+ab ema | normali zed cell [illegible] | 1d, 3d, 6d, 7d, 14d, 21d, each time point n=3 | Calibr ation | *Nalive* |
| 2 C | E2+ab ema | c-Myc | 1d, 3d, 6d, 7d, each time point n=3 | Calibr ation | *cMyc* |
| 2 D | E2+ab ema | RB1-pp | 1d, 3d, 6d, 7d, each time point n=3 | Calibr ation | *ppRb* |

| | | | | | |
|---|---|---|---|---|---|
| 3A | Alternating treatment | normalized cell number | 7d, E2+palbo(250nM)<br>14d, –E2<br>21d, E2+palbo(250nM) | Calibration | *Nalive* |
| 3B | Alternating treatment | normalized cell number | 7d, E2+palbo(500nM)<br>14d, E2+ICI(500nM)<br>21d, E2+palbo(250nM) | Calibration | *Nalive* |
| 3C | Alternating treatme | normalized cell number | 7d, E2+palbo(750nM)<br>14d, E2+ICI(500nM) | Prediction | *Nalive* |
| 3D | Alternating treatment | normalized cell number | 7d, E2+palbo(750nM)+ICI(500nM) | Prediction | *Nalive* |
| 4A | Alternating treatment | normalized cell number | 7d, E2+palbo(750nM)<br>14d, E2+ICI(750nM)<br>21d, E2+palbo(750nM)<br>28d, E2+ICI(750nM)<br>35d, E2+palbo(750nM)<br>42d, E2+ICI(750nM)<br>49d, E2+palbo(750nM) | Calibration | *Nalive* |
| 4A | E2+palbo<br>(750nM | normalized cell number | 7d, 14d, 21d, 28d, 35d, 42d, 49d, 56d, 63d, 70d, each time point n=3 | Calibration | *Nalive* |
| 4A | E2+ICI<br>(750nM | normalized cell | 7d, 14d, 21d, 28d, 35d, each time point n=3 | Calibration | *Nalive* |
| 4E | E2+palbo<br>(750nM) | cyclinD1 | 35d, 50d, each time point n=3 | Calibration | *cyclinD1+*<br>*cyclinD1cdk46+*<br>*cyclinD1cdk46p21+*<br>*cyclinD1Cdk46palo+*<br>[illegible] |
| 4E | Alternating treatme | cyclinD1 | 35d, E2+palbo(750nM)<br>70d, E2+ICI(750nM) | Calibration | Same as above |
| 4F | E2+palbo<br>(750nM) | Cdk4 | 35d, 50d, each time point n=3 | - | *cdk46+*<br>*cdk46palbo+*<br>*cdk46abema+*<br>*cyclinD1cdk46+*<br>*cyclinD1cdk46p21+* |
| 4F | Alternating treatme | Cdk4 | 35d, E2+palbo(750nM)<br>70d, E2+ICI(750nM) | - | Same as above |

| | | | | | |
|---|---|---|---|---|---|
| 4G | E2+palbo | Cdk6 | 35d, 50d, each time point n=3 | - | Same as above |
| 4G | Alternating | Cdk6 | 35d, E2+palbo(750nM) [illegible] | - | Same as above |
| 4H | E2+palbo | cyclinE | 35d, 50d, each time point n=3 | - | *cyclinE+cyclinEp21* |
| 4H | Alternating | cyclinE | 35d, E2+palbo(750nM) [illegible] | - | *cyclinE+cyclinEp21* |
| 4I | E2+palbo | Cdk2 | 35d, 50d, each time point n=3 | - | - |
| 4I | Alternating | Cdk2 | 35d, E2+palbo(750nM) [illegible] | - | - |
| 4J | E2+palbo (750nM) | cyclinD1 | 7d, E2+palbo(750nM)<br>14d, E2+ICI(750nM)<br>each time point n=3 | - | *cyclinD1+*<br>*cyclinD1cdk46+*<br>*cyclinD1cdk46p21+*<br>*cyclinD1Cdk46palo+* |
| 6J | E2+ICI [illegible] | normalized cell [illegible] | 17d, n=3 | Prediction | *Nalive* |
| 6J | E2+palbo [illegible] | normalized cell number | 17d, n=3 | Prediction | *Nalive* |
| 6J | E2+palbo [illegible] | normalized cell number | 17d, n=3 | Prediction | *Nalive* |
| 6J | E2+palbo [illegible] | normalized cell number | 17d, n=3 | Prediction | *Nalive* |
| 6K | E2+ICI [illegible] | normalized cell [illegible] | 17d, n=3 | Prediction | *Nalive* |
| 6K | E2+abema [illegible] | normalized cell number | 17d, n=3 | Prediction | *Nalive* |
| 6K | E2+abema [illegible] | normalized cell number | 17d, n=3 | Prediction | *Nalive* |
| S1 | –E2 | c-Myc | 1d, 3d, 6d, 7d, each time point n=3 | Calibration | *cMyc* |
| S1A | –E2 | cyclinD1 | 1d, 3d, 6d, 7d, each time point n=3 | Calibration | *cyclinD1+*<br>*cyclinD1cdk46+*<br>*cyclinD1cdk46p21+*<br>*cyclinD1Cdk46palo+* |
| S1 | –E2 | ERα | 1d, 3d, 6d, 7d, each time point n=3 | Calibration | *ER+E2ER+ ICIER* |
| S1 | –E2 | RB1-pp | 1d, 3d, 6d, 7d, each time point n=3 | Calibration | *ppRb* |

| | | | | | |
|---|---|---|---|---|---|
| S1 | –E2 | RB1 | 1d, 3d, 6d, 7d, each time point n=3 | Calibration | *Rb+pRb+ppRb* |
| S2 | E2+ICI (500nM [illegible] | c-Myc | 1d, 3d, 6d, 7d, each time point n=3 | Calibration | *cMyc* |
| S2 | E2+ICI (500nM) | cyclinD1 | 1d, 3d, 6d, 7d, each time point n=3 | Calibration | *cyclinD1+*<br>*cyclinD1cdk46+*<br>*cyclinD1cdk46p21+*<br>*cyclinD1Cdk46palo+* |
